# Mobile brain/body imaging of landmark-based navigation with high-density EEG

**DOI:** 10.1101/2021.01.13.426330

**Authors:** Alexandre Delaux, Jean-Baptiste de Saint Aubert, Stephen Ramanoël, Marcia Bécu, Lukas Gehrke, Marius Klug, Ricardo Chavarriaga, José-Alain Sahel, Klaus Gramann, Angelo Arleo

**Affiliations:** Sorbonne Universités, INSERM, CNRS, Institut de la Vision, Paris, France; Institute of Psychology and Ergonomics, Technische Universität Berlin, Berlin, Germany; Center for Neuroprosthetics, Ecole Polytechnique Fédérale de Lausanne, Geneva, Switzerland; Zurich University of Applied Sciences, ZHAW Datalab, Winterthur, Switzerland; CHNO des Quinze-Vingts, INSERM-DGOS CIC 1423, Paris, France; Fondation Ophtalmologique Rothschild, Paris, France; Department of Ophthalmology, The University of Pittsburgh School of Medicine, Pittsburgh, United States

**Keywords:** mobile EEG, virtual reality, ecological navigation, source reconstruction, retrosplenial complex (RSC)

## Abstract

Coupling behavioral measures and brain imaging in naturalistic, ecological conditions is key to comprehend the neural bases of spatial navigation. This highly-integrative function encompasses sensorimotor, cognitive, and executive processes that jointly mediate active exploration and spatial learning. However, most neuroimaging approaches in humans are based on static, motion constrained paradigms and they do not account for all these processes, in particular multisensory integration. Following the Mobile Brain/Body Imaging approach, we aimed to explore the cortical correlates of landmark-based navigation in actively behaving young adults, solving a Y-maze task in immersive virtual reality. EEG analysis identified a set of brain areas matching state-of-the-art brain imaging literature of landmark-based navigation. Spatial behavior in mobile conditions additionally involved sensorimotor areas related to motor execution and proprioception usually overlooked in static fMRI paradigms. Expectedly, we located a cortical source in or near the posterior cingulate, in line with the engagement of the retrosplenial complex in spatial reorientation. Consistent with its role in visuo-spatial processing and coding, we observed an alpha power desynchronization while participants gathered visual information. We also hypothesized behavior-dependent modulations of the cortical signal during navigation. Despite finding few differences between the encoding and retrieval phases of the task, we identified transient time-frequency patterns attributed, for instance, to attentional demand, as reflected in the alpha/gamma range, or memory workload in the delta/theta range. We confirmed that combining mobile high-density EEG and biometric measures can help unravel the brain structures and the neural modulations subtending ecological landmark-based navigation.

## Introduction

Spatial navigation requires active exploration, multisensory integration, as well as the encoding and long-term consolidation of internal models of the world (Arleo & Rondi-Reig, 2007; Wolbers & Hegarty, 2010; Epstein *et al*., 2017). Thus, the ability to navigate in space encompasses both perceptual and cognitive faculties (Spiers & Barry, 2015; Ekstrom *et al*., 2017). A large body of work has elucidated the neural bases of wayfinding behavior in both animals and humans, leading to a better understanding of the navigational system across multiple levels (Burgess, 2008; Epstein *et al*., 2017; Hardcastle *et al*., 2017; Poulter *et al*., 2018).

Most investigations of the brain network subtending human spatial navigation rely on functional magnetic resonance imaging (fMRI) (Taube *et al*., 2013; Epstein *et al*., 2017), due to its unmatched spatial resolution among non-invasive methods. However, this technique is not suited for testing participants in unconstrained motion conditions, which limits the study of neural processes involved during natural behavior (Zaitsev *et al*., 2015). Combining behaviometric and neurometric recordings in ecological (i.e., close to real, natural) conditions is key to modern cognitive neuroscience (Schaefer, 2014; Ladouce *et al*., 2019), in particular to study spatial cognition (Bécu *et al*., 2020a; Gehrke & Gramann, 2021; Miyakoshi *et al*., 2021). With relatively coarse spatial but fine temporal resolution, electroencephalography (EEG) offers a complementary tool for neuroimaging the brain during spatial behavior (Bischof & Boulanger, 2003; Baker & Holroyd, 2009; Lin *et al*., 2009; Plank *et al*., 2010; Lin *et al*., 2015). Although EEG does not prevent the participant’s motion per se, it is very sensitive to movement-related artifacts. Electrical potentials from muscle contractions (e.g., head movements, eye blinks or heartbeat, see Jung *et al*., 2000) generate strong artifactual signals that compromise the extraction of brain-related responses (i.e., reducing the signal-to-noise ratio). As a consequence, most EEG studies have constrained the mobility of participants in order to minimize motion-related artifacts (e.g., by making them sit in front of a screen and respond with finger taps only).

Recent technical developments have unlocked the possibility of using EEG brain imaging in a variety of ecological conditions (indoor walking: Luu *et al*., 2017a; Ladouce *et al*., 2019; Park & Donaldson, 2019; outdoor walking: Debener *et al*., 2012; Reiser *et al*., 2019; cycling: Zink *et al*., 2016; di Fronso *et al*., 2019; dual-tasking: Marcar *et al*., 2014; Bohle *et al*., 2019). By coupling EEG recordings with other biometric measures (e.g., body and eye movements), the Mobile Brain/Body Imaging (MoBI) approach gives access to unprecedented behavioral and neural data analysis (Makeig *et al*., 2009; Gramann *et al*., 2011, 2014; Ladouce *et al*., 2017). In addition, the MoBI paradigm has been successfully combined with fully immersive virtual reality (VR) protocols (Snider *et al*., 2013; Plank *et al*., 2015; Liang *et al*., 2018; Djebbara *et al*., 2019; Peterson & Ferris, 2019). Immersive VR allows near-naturalistic conditions to be reproduced, while controlling all environmental parameters (Parsons, 2015; Park *et al*., 2018; Starrett & Ekstrom, 2018; Diersch & Wolbers, 2019). The reliability of 3D-immersive VR enables the stimulation of visual, auditory, and proprioceptive modalities, while allowing the participant to actively explore and sense the virtual environment (Bohil *et al*., 2011; Kober *et al*., 2012). This continuous interplay between locomotion and multisensory perception is thought to be a key component of spatial cognition in near-natural conditions, as its absence leads to impaired performance in various spatial abilities (path integration: Chance *et al*., 1998; spatial updating: Klier & Angelaki, 2008; spatial reference frame computation: Gramann, 2013; spatial navigation and orientation: Taube *et al*., 2013; Ladouce *et al*., 2017; spatial memory: Holmes *et al*., 2018).

In the present study, we use the MoBI approach to combine high-density mobile EEG recordings and immersive VR in order to study spatial navigation in a three-arm maze (i.e., a Y-maze). Our primary aim is to provide a proof-of-concept in terms of EEG-grounded neural substrates of landmark-based navigation consistent with those found in similar fMRI paradigms (Iaria *et al*., 2003; Wolbers *et al*., 2004; Wolbers & Büchel, 2005; Konishi *et al*., 2013). We chose the Y-maze task because it offers a simple two-choice behavioral paradigm suitable to study landmark-based spatial navigation and to discriminate between allocentric (i.e., world-centered) and egocentric (i.e., self-centered) responses, as previously shown in animals (Barnes *et al*., 1980) and humans (Rodgers *et al*., 2012; Bécu *et al*., 2020b). Complementarily, a recent fMRI study of ours has investigated the brain activity of regions involved in visuo-spatial processing and navigation in a similar Y-maze task (Ramanoël, Durteste, *et al*., 2020). This offers the opportunity to comparatively validate the neural correlates emerged through static fMRI experiment against those found by mobile high-density EEG.

The neural substrates of landmark-based navigation form a network spanning medial temporal areas (e.g., hippocampus, para-hippocampal cortex) and medial parietal regions (Epstein & Vass, 2014), such as the functionally-defined retrosplenial complex (RSC) (Epstein, 2008). Here, we expect the RSC to play a role in mediating spatial orientation through the encoding and retrieval of visual landmarks (Spiers & Maguire, 2006; Auger *et al*., 2012, 2015; Marchette *et al*., 2015; Auger & Maguire, 2018; Julian *et al*., 2018). The RSC is indeed implicated in the translation between landmark-based representations in both egocentric and allocentric reference frames (Vann *et al*., 2009; Sulpizio *et al*., 2013; Marchette *et al*., 2014; Shine *et al*., 2016; Mitchell *et al*., 2018). Our hypothesis also encompasses the role of specific upstream, visual processing areas of the parieto-occipital region involved in active wayfinding behavior (Bonner & Epstein, 2017; Patai & Spiers, 2017). In addition, our paradigm accounts for the role of downstream, higher-order cognitive functions necessary for path evaluation, covering a frontoparietal network (including prefrontal areas, Spiers & Gilbert, 2015; Epstein *et al*., 2017) that codes for overarching mechanisms such as spatial attention and spatial working memory (Cona & Scarpazza, 2019).

Mobile brain imaging protocols also engage locomotion control processes, in which motor areas in the frontal lobe and somatosensory areas in the parietal lobe are typically involved (Gwin *et al*., 2010; Seeber *et al*., 2014; Roeder *et al*., 2018; see Delval *et al*., 2020 for a recent review). Furthermore, the integration of vestibular and proprioceptive cues made possible by mobile EEG paradigms is likely to influence the observed neural correlates of spatial orientation (Ehinger *et al*., 2014; Gramann *et al*., 2018) and attention (Ladouce *et al*., 2019).

Finally, given the high temporal resolution of EEG, we aim at characterizing how the activity of the structures engaged in active, multimodal landmark-based navigation is modulated by behavioral events, related to either action planning (e.g., observation of the environment, physical rotation to complement mental perspective taking) or action execution (e.g., walking, maintaining balance). We also aim at exploring the differential engagement of brain regions involved in the encoding (learning condition) and the retrieval (control and probe conditions) phases of the task (RSC is implicated in both; (Epstein & Vass, 2014; Burles *et al*., 2018; Mitchell *et al*., 2018).

The purpose of this study is thus to explore the cortical correlates of landmark-based navigation in mobile participants. We first hypothesize that the analysis of the EEG signal will retrieve the above-mentioned brain structures known to be engaged during active spatial navigation based on visual cues. We then expect behavioral events to modulate features of the recorded EEG data, identifiable as transient time-frequency patterns in the involved brain areas, and to interpret them with respect to spatial cognition and locomotion control literature. Finally, we expect to find significant differences in these patterns across the phases of the task, contrasting the cognitive mechanisms involved in context-dependent task solving. We aim to investigate and interpret their condition-specificity and their temporality. Under such considerations, this work can help towards a better understanding of context-specific neural signatures of landmark-based navigation.

## Methods

### Participants

Seventeen healthy adults (range, 21-35 years old, M = 26,82, SD = 4,85, 10 females) participated to this study. Fifteen were righthanded and two lefthanded. All participants had normal (or corrected to normal) vision and no history of neurological disease. In one recording session, there were abnormalities (discontinuities, absence of events) in the motion capture signal. Thus, we removed one participant from the analysis. The experimental procedures were approved by the local ethics committee (GR_12_20190513, Institute of Psychology & Ergonomics, *Technische Universität Berlin*, Germany) and all participants signed a written informed consent, in accordance with the Declaration of Helsinki. All participants answered a discomfort questionnaire at the end of the experiment, adapted from the simulation sickness questionnaire of Kennedy *et al*., (1993), which can be found in Supplementary Methods 1. We gave the instructions in English and all participants reported a good understanding of the English language. Each participant received a compensation of either 10€/h or course credits.

### EEG system

The EEG system (Fig. 1a) consisted of 128 active wet electrodes (actiCAP slim, Brain Products, Gilching, Germany) mounted on an elastic cap with an equidistant layout (EASYCAP, Herrsching, Germany). The impedance of a majority of the channels was below 25 kΩ (9.5% of the electrodes had an impedance above 25 kΩ). Two electrodes placed below the participant’s eyes recorded electro-oculograms (EOG). An additional electrode located closest to the standard position F3 (10-20 international system) provided the reference for all other electrodes. The EEG recordings occurred at a sampling rate of 1000 Hz. The raw EEG signal was streamed wirelessly (BrainAmp Move System, Brain Products, Gilching, Germany) and it was recorded continuously for the entire duration of the experiment.

**Figure 1.**
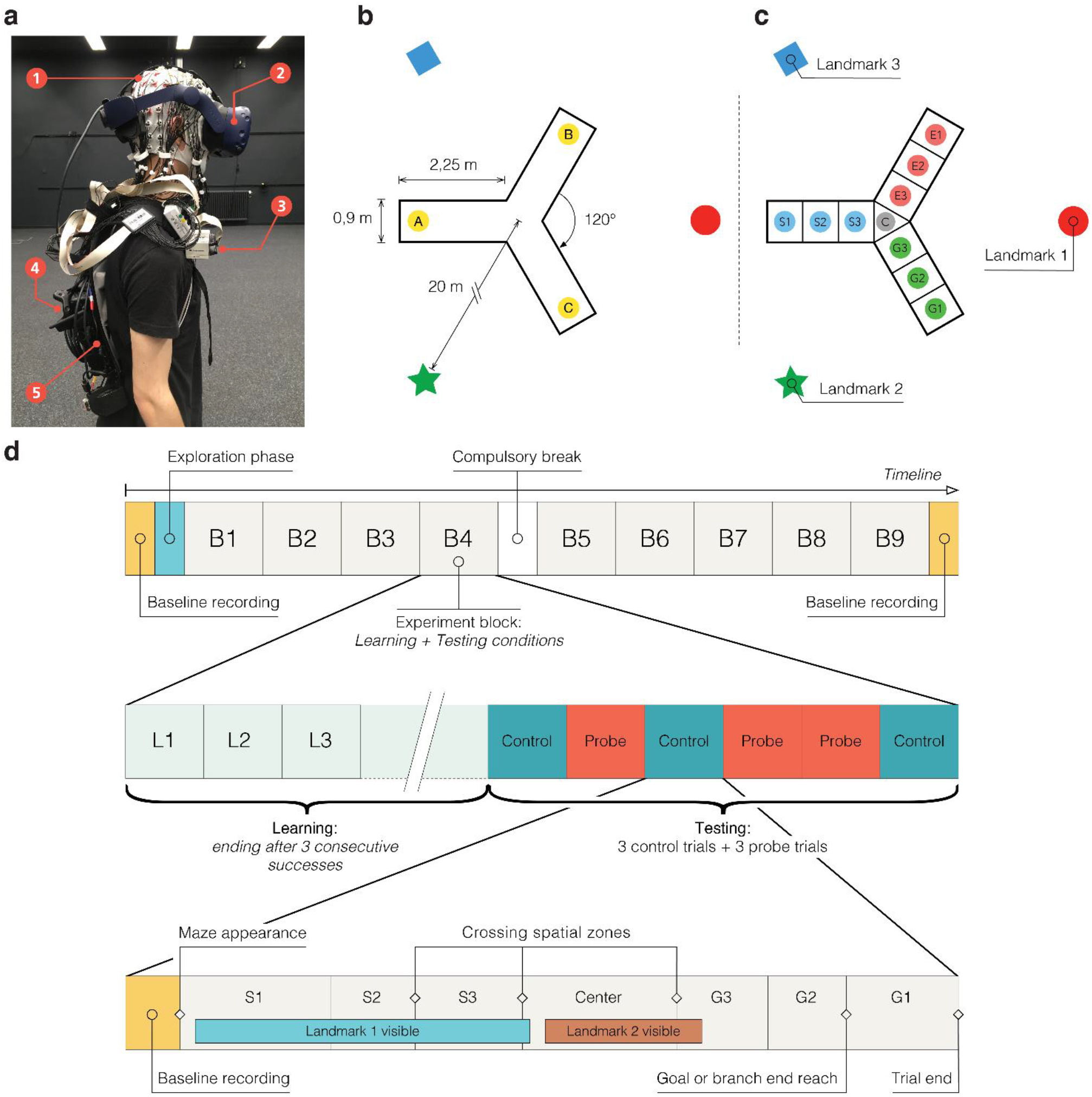
Virtual environment, setup, and timeline of the experiment. **(a)** Details of participant’s equipment. (1) EEG cap (128 channels) - (2) VR Head-mounted display (VIVE Pro) - (3) Wifi transmitter for EEG data (Move system) - (4) Additional motion capture tracker (VIVE tracker) - (5) Backpack computer running the virtual environment (Zotac PC). **(b)** Virtual environment. Participants explored a virtual equilateral Y-maze. In the learning condition, they always started in the same arm (e.g., A) and they had to find a hidden goal, always placed in the same location (e.g., C). In the testing conditions, the environment and goal location stayed the same but the participant would start from either the same position (A) in control trials or the third arm (B) in probe trials. **(c)** Spatial discretization of the environment (example for a learning trial). We delimited 10 areas in the maze: ‘S’ stands for starting arm, ‘C’ for center, ‘E’ for error arm, and ‘G’ for goal arm. In the text, when referring to the arm chosen by the participant (either ‘E’ or ‘G’), we use the letter ‘F’ standing for finish arm. These labels are condition-dependent (different in the probe condition). The names of the landmark depend on the location of starting arm in the learning condition and goal arm. These names are block dependent. **(d)** General timeline of the experiment. The first row represents the general succession of conditions in the experiment. The second row shows an example of the sequence of trials in an experimental block. The third row illustrates the structure of a trial, including a possible course of events: progress across spatial sections and visibility of landmarks depending on participant’s head movements. We provide a video of a participant performing the task, along with the reconstruction of the tracker positions, in Supplementary Media.

### Virtual Y-maze and motion tracking

The virtual maze consisted of an equilateral Y-maze (3-armed maze) with three distal landmarks placed outside the maze, 20 m away from the center and visible above the walls (Fig. 1b). The landmarks were abstract geometric shapes (e.g., square, circle, star). The wall texture and the light were homogeneous and non-informative. Each arm of the maze was 90 cm wide and 225 cm long. For the sake of analysis, the maze was discretized into 10 zones (3 evenly divided zones per arm and one for the maze center, Fig. 1c). These zones were not visible to the participant. Crossing between zones was recorded online without influencing the task flow.

We designed the virtual Y-maze by using the Unity3D game engine (Unity Technologies, San Francisco, California, USA, version 2017.1.1f1 for Windows), and we rendered it using an HTC Vive Pro head-mounted display (HTC Corporation, Taoyuan, Taïwan) with a 90 Hz refresh rate (2 times AMOLED 3.5” 1440×1600 pixels, 615 ppi and 110° nominal field of view). The HTC was connected to a VR capable backpack computer (Zotac PC, Intel 7th Gen Kaby Lake processor, GeForce GTX 1060 graphics, 32GB DDR4-2400 memory support, Windows 10 OS, ZOTAC Technology Limited, Fo Tan, Hong Kong) running on batteries and controlled remotely (Fig. 1a). An integrated HTC Lighthouse motion tracking system (four cameras, 90 Hz sampling rate, covering an 8×12 meters area) enabled the recording of the participant’s head by tracking the HTC Vive Pro head-mounted display. It also enabled the tracking of the torso movements via an additional HTC Vive Tracker placed on the participant’s backpack. We virtually translated the position of this tracker to better reflect the real position of the participant’s torso by considering his or her body measurements. The torso tracker was also used to trigger spatial events (e.g., reaching the goal, crossing spatial section boundaries). The height of the maze walls and the altitude of landmarks were adjusted to the participant’s height (based on the head tracker) to provide each participant with the same visual experience. Each participant wore earphones playing a continuous white noise, to avoid auditory cues from the external world. During the disorientation periods (see protocol), relaxing music replaced the white noise. One experimenter gave instructions through the earphones, while monitoring the experiment from a control room. The participant could answer through an integrated microphone. He/she was instructed to refrain from talking while performing the experiment to limit artifacts in the recorded EEG signal. Another experimenter stayed with the participant inside the experimental room to help with potential technical issues and conduct the disorientation, avoiding any interaction with participants during the task. The EEG signal, motion capture, and all trigger events were recorded and synchronized using the Lab Streaming Layer software (Kothe, 2014).

### Experimental protocol

An entire experimental session lasted 3 hours on average and it included preparing the participant with the EEG and VR equipment and running the experimental protocol. The immersion time in VR was between 60 and 90 minutes.

#### Free exploration phase

Before starting the actual task, the participant explored the Y-maze for 3 minutes, starting at the center of the maze. He/she was instructed to inspect all details of the environment and to keep walking until the time elapsed. The purpose of this phase was to familiarize the participant to the VR system and the Y-maze environment (including the constellation of landmarks).

#### Navigation task

The navigation task included a learning condition and a testing condition. During learning, the participant began each trial from the *starting* arm (e.g., location A in Fig. 1b) and he/she had to find the direct route to a hidden target at the end of the *goal* arm (e.g., location C in Fig. 1b). Upon reaching the goal, a reward materialized in front of the participant (3D object on a small pillar representing, for instance, a treasure chest) to indicate the correct location and the end of the current trial. The learning period lasted until the participant reached the goal directly, without entering the other arm, three times in a row. Before each trial, we disoriented the participant to ensure that he/she would not rely on previous trials or the physical world to retrieve his/her position and orientation. To disorient the participant, the experimenter simply walked him/her around for a few seconds with both eyes closed (and the head-mounted display showing a black screen). The testing condition included 6 trials: 3 *control* trials and 3 *probe* trials, ordered pseudo-randomly (always starting with a control, but never with 3 control trials in a row). In the control trials, the participant started from the same arm as in the learning condition (e.g., location A in Fig. 1b). In the probe trials, he/she started from the third arm (e.g., location B in Fig. 1b). Before starting a new trial (either control or probe), the participant was always disoriented. Then, he/she had to navigate to the arm where he/she expected to find the goal and stop there (without receiving any reward signal). If the participant went to the incorrect arm, it was considered as an *error*. We present a single trial example of one participant performing the task and we illustrate the motion tracking in the virtual environment in Supplementary Video 1.

#### Block repetitions

The sequence “learning condition + testing condition” formed an *experimental block*. Each participant performed 9 experimental blocks (Fig. 1d). In order to foster a feeling of novelty across block repetitions, we varied several environmental properties at the beginning of each block: wall texture (e.g., brick, wood, etc.), goal location (i.e., in the right or left arm, relative to the starting arm), reward type (e.g., treasure chest, presents, etc.), as well as the shape (e.g., circle, square, triangle, etc.) and color of landmarks. When changing the environment between blocks, we kept the maze layout and landmark locations identical. The sequence of blocks was identical for all participants, who had to take a compulsory break after the fourth block (Fig. 1d). In addition, after the sixth or seventh block, a break was introduced when requested by the participant.

#### EEG baseline recordings

Both before the free exploration period and after the 9^th^ block, the participant had to stand for 3 minutes with his/her eyes opened in a dark environment. This served to constitute a general baseline for brain activity. Similarly, we recorded the EEG baseline signal (in the dark for a random duration of 2 to 4 seconds) before each trial (Fig. 1d, bottom). Besides providing a baseline EEG activity specific to each trial, this also allowed the starting trial time (i.e., the appearance time of the maze) to be randomized, thus avoiding any anticipation by the participant.

### Behavioral analysis

All analyses were done with MATLAB (R2017a and R2019a; The MathWorks Inc., Natick, Massachusetts, USA), using custom scripts based on the EEGLAB toolbox version 14.1.0b (Delorme & Makeig, 2004), the MoBILAB (Ojeda *et al*., 2014) open source toolbox, and the BeMoBIL pipeline (Klug *et al*., 2018).

#### Motion capture processing

A set of MoBILAB’s adapted functions enabled the preprocessing of motion capture data. The rigid body measurements from each tracker consisted of (x, y, z) triplets for the position and quaternion quadruplets for the orientation. After the application of a 6 Hz zero-lag lowpass finite impulse response filter, we computed the first time derivative for position of the torso tracker for walking speed extraction and we transformed the orientation data into axis/angle representations. An EEGLAB dataset allowed all preprocessed, synchronized data to be collected and split into different streams (EEG, Motion Capture) to facilitate EEG-specific analysis based on motion markers.

#### Allocentric and egocentric groups

Probe trials served to distinguish between allocentric and egocentric responses by making the participants start from a different arm than the one used in the learning period. We assigned a participant to the *allocentric group* if he/she reached the goal location in the majority of probe trials (i.e., presumably, by using the landmark array to self-localize and plan his/her trajectory). Conversely, we assigned a participant to the *egocentric group* when he/she reached the error arm in the majority of probe trials (i.e., by merely repeating the right- or left-turn as memorized during the learning period).

#### Time to goal

We assessed the efficacy of the navigation behavior by measuring the “time to goal”, defined as the time required for the participant to finish a trial (equivalent to the “escape latency” in a Morris Water Maze). In learning trials, it corresponded to the time to reach the goal zone and trigger the reward. In test trials, it corresponded to the time to reach the believed goal location in the chosen arm (i.e., entering the G1 or E1 zone in Fig. 1c).

#### Horizontal head rotations (relative heading)

The participant’s heading was taken as the angle formed by his/her head orientation in the horizontal plane with respect to its torso orientation, aligned with the participant’s sagittal plane. After extracting the head and torso forward vectors from each tracker, computing the signed angle between those vectors’ projections in the horizontal plane provided the heading value.

#### Walking speed

The forward velocity component of the torso tracker provided the participant’s walking information. For each trial, we computed the mean and standard deviation (SD) of the forward velocity, and their average for each participant. To evaluate movement onsets and offsets, we compared motion data recorded during the trials against those recorded during the short baseline period before each trial, considered as a reliable resting state for movements. Movement transitions (onsets and offsets) were based on a participant-specific threshold, equal to the resting state mean plus 3 times the resting state SD. The excluded movement periods were those lasting less than 250 ms and during which motion did not reach another participant-specific threshold, equal to the resting state mean plus 5 times the resting state SD.

#### Landmark visibility

For analysis purposes, we named the three landmarks as Landmark 1, Landmark 2, and Landmark 3 (Fig. 1c) and we tracked their visibility within the displayed scene. The participant had a horizontal field of view of 110° and a vertical field of view of 60°. Whenever a landmark appeared in the viewing frustum^1^ (i.e., in the region of virtual space displayed on the screen) and it was not occluded by any wall, it was considered as visible by the participant. Given the restrained horizontal field of view and the configuration of the VR environment, perceiving more than one landmark at the same time was unlikely.

#### Zone-based behavioral analysis

The maze discretization (Fig. 1c) provided a coherent basis for analyses across trials and participants. To ensure consistency in the comparison between trials, we selected those trials where the participant followed a straightforward pattern (zone-crossing sequence: S1 ➔ S2 ➔ S3 ➔ C ➔ G3 ➔ G2 ➔ G1). We thus discarded all trials in which the participant went backward while navigating (e.g., during learning, when his/her first choice was towards the error arm, and he/she had to come back to the center in order to go towards the goal arm). To further ensure homogeneity, we also excluded those trials in which the time to goal was unusually long (i.e., by computing the outliers of the time to goal distribution across all participants). These selection criteria kept 1289 (out of a total of 1394) trials for analysis (see Supp. Table 1 for details about the distribution of trials across participants). Finally, we computed offline an additional event corresponding to the first walking onset of the participant in S1 (see ***Walking speed*** paragraph above for movement detection) which was inserted in the delimiting sequence of zone-crossing events. For the sake of simplicity, we used the notations “staticS1” for the period preceding this event, and “mobileS1” for the one that follows, before the participant enters S2. Hence, the complete sequence for each trial was, e.g., staticS1 ➔ mobileS1 ➔ S2 ➔ S3 ➔ C ➔ G3 ➔ G2 ➔ G1.

#### Motion capture statistics

The above zone-based discretization framed the analysis of the motion capture metrics mentioned above: walking speed, standard deviation of horizontal head rotations, and landmark visibility. For each trial, we first averaged the value of each motion variable over the period between two events of the zone sequence. Then, for each participant, we averaged these values across trials of the same condition.

To better characterize the participants’ behavior in the maze, we investigated how these metrics would depend on the condition, the spatial zone, the landmark (for landmark visibility only), and the different combinations of those factors. Concerning walking speed and standard deviation of horizontal head rotations, we tested the hypothesis that participants would walk slower and make larger head movements in specific zones of the maze related to the challenge posed by the experimental condition (e.g., taking information in S1 during Learning and stopping in C to look at the constellation during Probe). Concerning landmark visibility, we tested the hypothesis that participants would make a differential use of the 3 landmarks (i.e., preference for one or two) and that attendance to a landmark would depend on the condition and the location of the participant in the maze (e.g., realignment with a preferred landmark at the center specific to Probe condition). We used fixed model between factors analyses of variance (ANOVAs; balanced design) to assess differences and interactions between conditions, zones and landmarks in those dependent variables. Specifically, for the landmark visibility, we used a 3-way ANOVA with the factors: *condition* (Learning, Control, Probe), *landmark* (Landmark 1, Landmark 2, Landmark 3) and *zone* (e.g., staticS1, mobileS1, S2, S3, Center, F3, F2). Note that ‘F’, standing here for ‘finish’ arm, can be either G for ‘goal’ or E for ‘error’ as used in Fig. 1c, depending on the trial outcome. For the walking speed and the standard deviation of horizontal head rotations, we used a 2-way ANOVA with the factors *condition* and *zone*. The alpha level for significance was set to 0.01 (more conservative level taking into account that we are computing three simultaneous ANOVAs on the same dataset). When a significant main effect or interaction was found, we used pairwise t-tests (with Tukey’s honest significant difference criterion method for multiple comparison correction) to unravel individual differences between factor or interaction terms.

### EEG data analysis

Figure 2 shows the outline of the data preprocessing and analysis steps.

**Figure 2.**
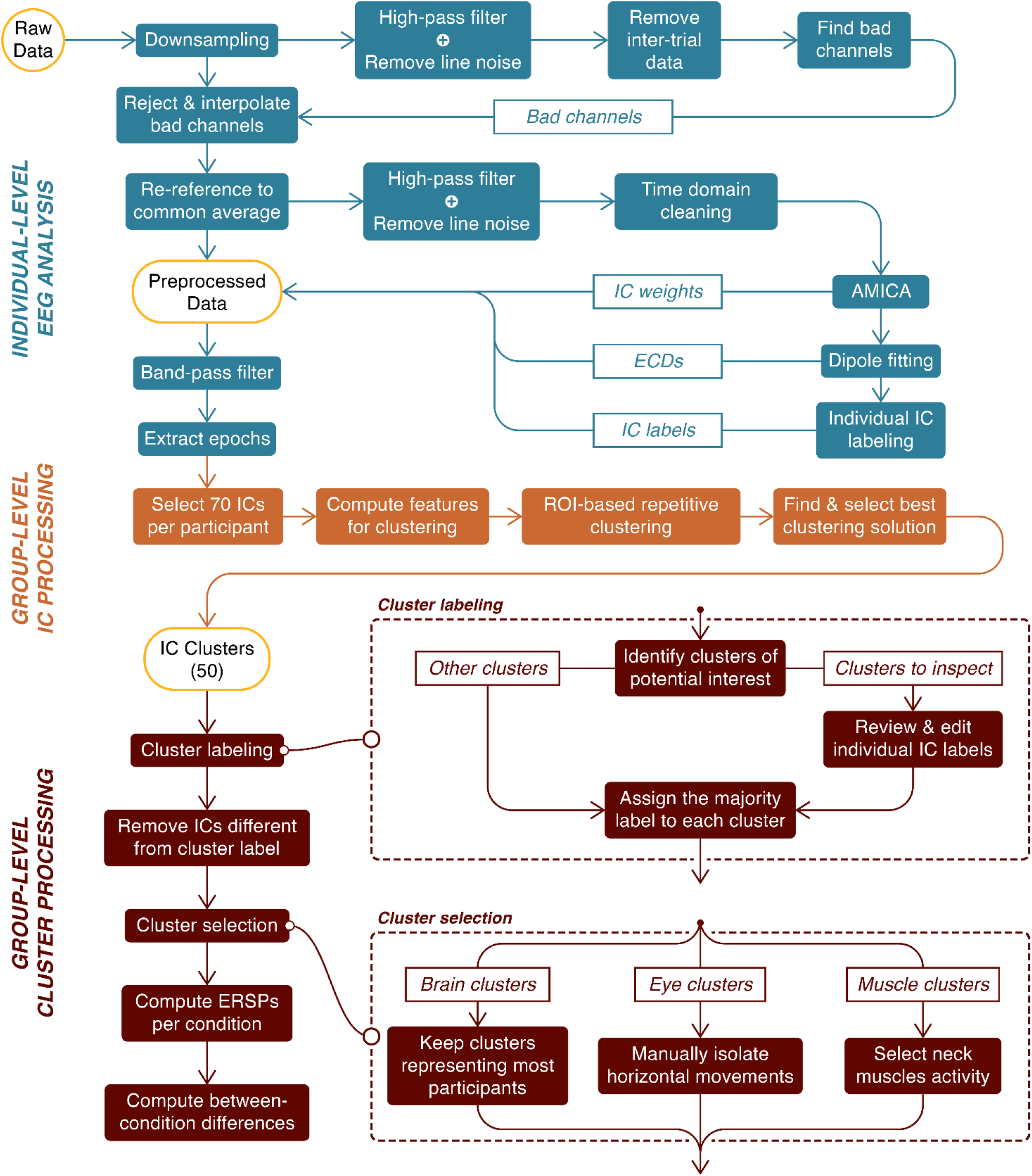
Flowchart of the EEG processing pipeline. We first preprocessed EEG data at the individual level (in blue) and, in particular, decomposed the channel data into independent components (ICs) with an adaptative mixture ICA (AMICA) algorithm. We then selected 70 ICs per participant for the clustering procedure (in orange). Finally, we labeled and selected the clusters of interest for an ERSP analysis per condition (in brown). The “Cluster selection” process is described in the “EEG cluster analysis” section of the Results.

#### Individual EEG analysis

##### Processing

We used the BeMoBIL pipeline to preprocess and clean the EEG data (Klug *et al*., 2018). This pipeline is fully automated and designed to improve signal to noise ratio (SNR) in large-scale mobile EEG datasets, which ensures full replicability of the procedure. We first downsampled the data to 250 Hz, applied a 1 Hz high-pass filter to suppress slow drifts in EEG data (zero-phase Hamming windowed finite impulse response filter with 0.5 Hz cut-off frequency and 1 Hz transition bandwidth), and removed spectral peaks at 50 Hz and 90 Hz, corresponding to power line frequency and VIVE headset refreshing rate, respectively (implemented by the *cleanLineNoise* function from the PREP pipeline, Bigdely-Shamlo *et al*., 2015). We identified noisy channels with automated rejection functions, setting parameters numerical values according to default recommendations from Bigdely-Shamlo *et al*., (2015). We then reconstructed the removed channels by spherical interpolation of neighboring channels and applied re-referencing to the common average. In a subsequent time-domain cleaning, we detected and removed segments with noisy data. We present more details on the implementation of the cleaning steps in Supplementary Methods 2.

On the cleaned dataset, we performed an independent component analysis (ICA) using an adaptive mixture independent component analysis (AMICA) algorithm (Palmer *et al*., 2008), preceded by a principal component analysis reduction to the remaining rank of the dataset taking into account the number of channels interpolated and the re-referencing to the common average. For each independent component (IC), we computed an equivalent current dipole model (ECD) with the DIPFIT plugin for EEGLAB (version 3.0) (Oostenveld & Oostendorp, 2002). For this purpose, we used a common electrode location file obtained from the average of previous measures on participants wearing the same cap. We co-registered this file with a boundary element head model based on the MNI brain (Montreal Neurological Institute, MNI, Montreal, QC, Canada) to estimate dipole location. In this article, the spatial origin of an IC is approximated with the location of its associated dipole.

We opted for the BeMoBIL pipeline after comparing it against the APP pipeline (da Cruz *et al*., 2018), which proved to be less robust for our dataset. We based this conclusion on different metrics, by evaluating each artifactual detection step (number of channels removed, proportion of time samples excluded) and by assessing the performance of the subsequent ICA (mutual information reduction and remaining pairwise mutual information, Delorme *et al*., 2012). In particular, the BeMoBIL pipeline proved to be more stable and conservative than the APP pipeline (rejecting more artifactual channels and noisy temporal segments, both more consistently across participants). We detail the comparison and its results in Supplementary Methods 3 and Supplementary Figure 4, respectively.

##### Individual IC labeling

We used the ICLabel algorithm (version 1.1, Pion-Tonachini *et al*., 2019) with the ‘default’ option to give an automatic class prediction for each IC. The model supporting this algorithm considers 7 classes: (1) Brain, (2) Muscle, (3) Eye, (4) Heart, (5) Line Noise, (6) Channel Noise and (7) Other. The prediction takes the form of a compositional label: a percentage vector expressing the likelihood of the IC to belong to each of the considered classes. Then it compares each percentage to a class-specific threshold to form the IC label. We used the threshold vector reported by Pion-Tonachini *et al*. (2019) for optimizing the testing accuracy. Considering the recentness of this algorithm and the fact it has never been validated on mobile EEG data, we refined the labeling process to increase its conservativeness on Brain ICs. After the initial categorization by the algorithm, we automatically examined the ECD of ICs passing the Brain threshold and we rejected all ICs whose ECD was either located outside brain volume or exhibiting residual variance over 15% (commonly accepted threshold for dipolarity, see Delorme *et al*., 2012) and we put them in the “Other” class. Residual variance quantifies the quality of the fit between the actual topographic activation map and the estimated dipole projection on the scalp. Among the remaining ones, we distinguished 2 cases: (1) if the IC label was uniquely “Brain”, we automatically accepted it; (2) if the IC label was hybrid (multiple classes above threshold), we manually inspected the IC properties to assign the label ourselves according to the ICLabel guidelines (https://labeling.ucsd.edu/tutorial/labels – an example can be found in Supp. Figure 3). To all ICs below brain threshold, we assigned unique labels based on their highest percentage class.

#### Group level EEG analysis

In order to retain maximal information for further processing, for each participant we copied the ICA results (decomposition weights, dipole locations and labels) back to the continuous version of the dataset (i.e., the dataset before time domain cleaning in the BeMoBIL pipeline). We first band-pass filtered the data between 1 Hz (zero-phase Hamming windowed finite impulse response high-pass filter with 0.5 Hz cut-off frequency and 1 Hz transition bandwidth) and 40 Hz (zero-phase Hamming windowed finite impulse response low-pass filter with 45 Hz cut-off frequency and 10 Hz transition bandwidth). We then epoched each dataset into trials, starting at the beginning of the baseline period and ending at the time of trial completion. For each IC and each trial, we computed the trial spectrum using the pwelch method (1s Hamming windows with 50% overlap for power spectral density estimation). We baselined the spectrum with the average IC spectrum over all baseline periods using a gain model.

We additionally computed single trial spectrograms using the *newtimef* function of EEGLAB (1 to 40 Hz in linear scale, using a wavelet transformation with 3 cycles for the lowest frequency and a linear increase with frequency of 0.5 cycles). Using a gain model, we individually normalized each trial with its average over time (Grandchamp & Delorme, 2011). Separately for each participant, we calculated a common baseline from the average of trial baseline periods (condition-specific) and we subsequently corrected each trial with the baseline corresponding to its experimental condition (gain model). At the end, power data were log-transformed and expressed in decibels. To enable trial comparability, these event-related spectral perturbations (ERSPs) were time-warped based on the same sequence of events as for the zone-based analysis.

##### Component clustering

To allow for a group-level comparison of EEG data at the source level (ICs), we selected the 70 first ICs outputted by the AMICA algorithm, which corresponded to the ICs explaining most of the variance in the dataset (Gramann *et al*., 2018). This ensured the conservation of 90.6±1.8% (mean ± standard error) of the total variance in the dataset while greatly reducing computational cost and mainly excluding ICs with uncategorizable patterns. We conducted this selection independently of the class label for each IC. We applied the repetitive clustering region of interest (ROI) driven approach described in Gramann *et al*. (2018). We tested multiple sets of parameters to opt for the most robust approach and we present here the selected one (the detailed procedure for this comparison can be found in Supp. Methods 4 and its results in Supp. Table 3). We represented each IC with a 10-dimensional feature vector based on the scalp topography (weight=1), mean log spectrum (weight=1), grand average ERSP (weight=3), and ECD location (weight=6). We compressed the IC measures to the 10 most distinctive features using PCA. We repeated the clustering 10000 times to ensure replicability. According to the results from parameters comparison (see Supp. Methods 4), we set the total number of clusters to 50 and the threshold for outlier detection to 3 SD in the k-means algorithm. This number of clusters was chosen inferior to the number of ICs per participant to favor the analysis of clusters potentially regrouping ICs from a larger share of participants and therefore more representative of our population. We defined [0, −55, 15] as the coordinates for our ROI, a position in the anatomical region corresponding to the retrosplenial cortex (BA29/BA30). We set the first coordinate (x) to 0 because we did not have any expectation for lateralization. Coordinates are expressed in MNI format. We scored the clustering solutions following the procedure described in Gramann *et al*. (2018). For each of the 10000 clustering solutions, we first identified the cluster whose centroid was closest to the target ROI. Then, we inspected it using 6 metrics representative of the important properties this cluster should fulfill (see Supp. Table 3 detailing the comparison procedure results). In order to combine these metrics into a single score using a weighted sum (same weights used to choose the best of the 10000 solutions), we linearly scaled each metric value between 0 and 1. We eventually ranked the clustering solutions according to their score and selected the highest rank solution for the subsequent data analysis.

##### Cluster labeling

We then inspected the 50 clusters given by the selected clustering solution. We first used the individual IC class labels to compute the proportion of each class in the clusters. Since the clustering algorithm was blind to the individual class labels, most clusters contained ICs with heterogeneous labels. Bearing in mind that the ICLabel algorithm has not been validated on mobile EEG data yet, we suspected that the observed heterogeneity could, to a certain extent, owe to individual labeling mistakes. We therefore performed a manual check (identical to the hybrid case in the *Individual IC labeling* section above) of individual IC labels in specific clusters exhibiting a potential interest for the analysis. These clusters were those with at least 20% of Brain label, those with at least 50% of Eye label and those located in the neck region with at least 50% of Muscle label. Indeed, both eye and muscle activity are inherent to the nature of the mobile EEG recordings and their analysis can inform us on participants’ behavior (Gramann *et al*., 2014), similarly to horizontal head rotations and landmark visibility variables, with a finer temporal resolution. We finally labeled every cluster from their most represented class after correction, only when this proportion was above 50%. Eventually, within each of the labeled clusters, we removed the ICs whose label did not coincide with the cluster label.

##### Clusters analysis

We computed single-trial ERSPs as for the clustering procedure. To get the cluster-level ERSP, we took the arithmetic mean of the power data first at the IC level (including the baseline correction), then at the participant level, and finally at the behavioral group level. At the end of these operations, we log-transformed the power data to present results in decibels. We performed statistical analysis comparing ERSP activity between trial type (learning, control, probe), using a non-parametric paired permutation test based on maximum cluster-level statistic (Maris & Oostenveld, 2007) with 1000 permutations. For each permutation, we computed the F-value for each ‘pixel’ (representing spectral power at a given time-frequency pair) with an 1×3 ANOVA. Since the ANOVA test is parametric, we used log-transformed data for statistical analysis as ERSP sample distribution has a better accordance with gaussian distribution in that space (Grandchamp & Delorme, 2011). We selected samples with *F*-value above 95^th^ quantile of the cumulative *F*-distribution and clustered them by neighborhood. The cluster level *F*-value was the cumulative *F*-value of all samples in the cluster. We then formed the distribution of observed maximum clustered *F*-values across permutations to compute the Monte Carlo p-value for the original repartition. As a post-hoc test, we repeated the same analysis for each pair of conditions, with *t*-values instead of *F*-values and two-tailed t-test instead of ANOVA. We finally plotted ERSP differences only showing samples significant for both the 3 conditions permutation test and the inspected pairwise permutation test. The significance level was *p* < 0.05 for all tests in this case.

## Results

### Behavioral results

#### Goal-oriented navigation performance

During control trials, all participants successfully solved the Y-maze task by consistently choosing the goal arm (Supp. Fig. 1a, left). During the probe trials, 14 participants navigated to the correct goal arm (i.e., allocentric response), whereas 2 participants went to the error arm (i.e., egocentric response; Supp. Fig. 1a, right).

In terms of time to goal, all participants learned rapidly to locate and navigate to the goal position: after the first learning trial, in which goal finding was merely random, the mean time to goal of the allocentric group plateaued at around 6 s (Supp. Fig. 1b, left). During control trials, the mean time to goal of allocentric participants remained constant and identical to the plateau reached at the end of the learning condition (Supp. Fig. 1b, middle). In the probe trials, the mean time to goal of the allocentric group increased slightly by ~1s as compared to the control condition (Supp. Fig. 1b, right). Overall, the interindividual variability remained very low, reflecting the simplicity of the navigation task.

#### Spatial behavior across conditions and maze zones

We sought to characterize the exploratory behavior as a function of the protocol conditions (Condition factor) as well as of the zones in the Y-maze (Zone factor, see Fig. 1c). Hereafter, only the analyses on the allocentric group are presented as only two participants adopted an egocentric behavior (expectedly, Bécu *et al*., 2020b; see Supp. Fig. 2 for the individual behavior of egocentric participants).

##### Horizontal head rotations

First, we assessed the searching behavior by quantifying the horizontal head rotations variability (Figs. 3a,b). We did not observe any effect of Condition (*F*(2;273) = 2.69, *p* = 0.069), whereas we found a significant effect of the Zone on horizontal head rotations variability (*F*(6;273) = 8.99, *p* < 0.00001). Post-hoc analysis indicated that horizontal head rotations variability was higher at the beginning of the trajectory in comparison to the center of the maze (staticS1 vs. C, t(2) = 3.67, p < 0.01; mobileS1 vs. C, t(2) = 5.5, p < 0.00001). There was no interaction between Condition and Zone for this metric (*F*(12;273) = 0.13, *p* = 0.99).

**Figure 3.**
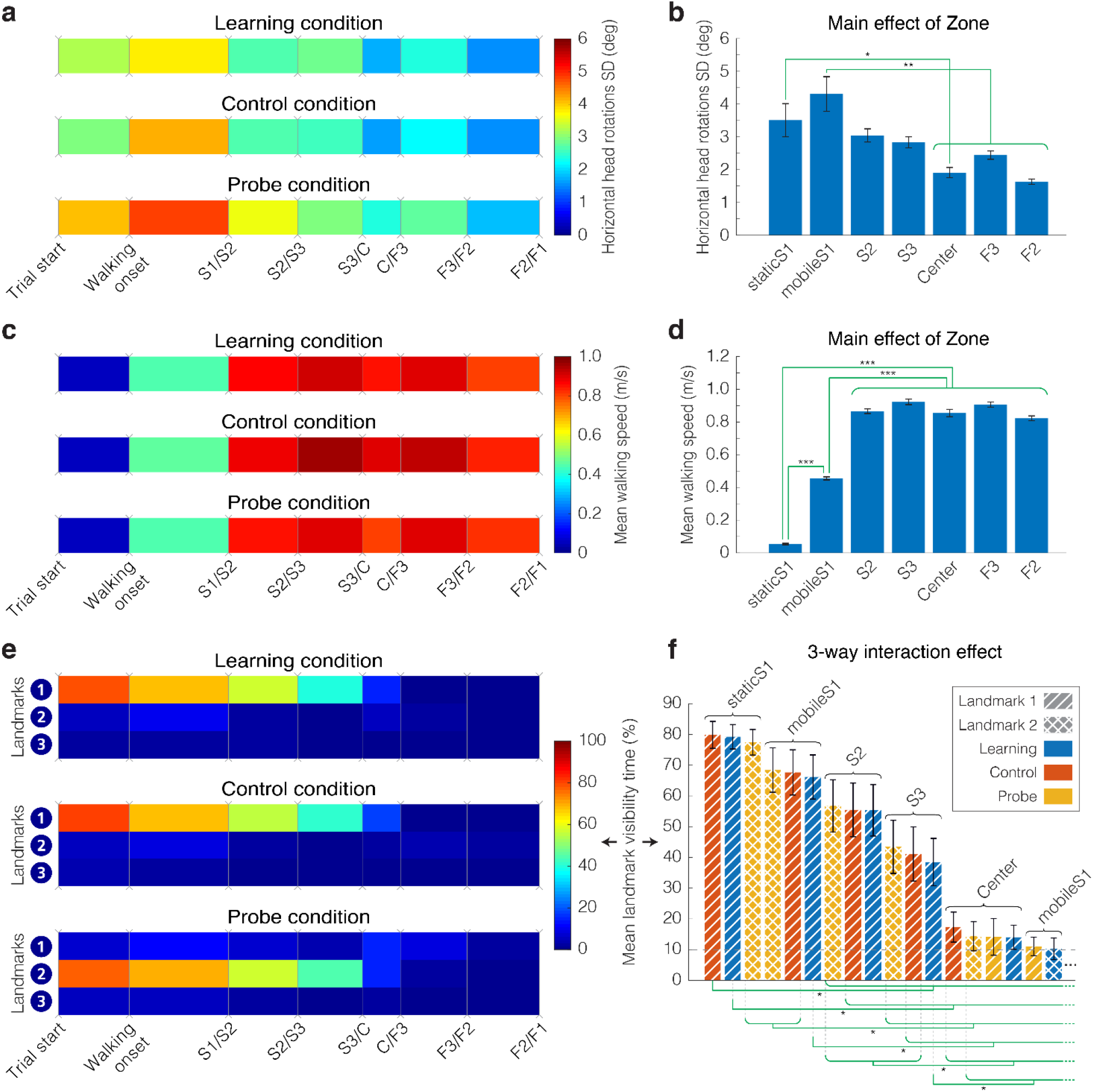
Behavioral metrics – Walking speed, horizontal head rotations variability, and landmark visibility for the allocentric group. **(a)** Average standard deviation of horizontal head rotations, computed from the difference between head and torso orientation. (b) Main effect of Zone on horizontal head rotations variability F(6;273) = 8.99, p < 0.00001. (c) Average instantaneous walking speed. (d) Main effect of Zone on walking speed F(6;273) = 472.15, p < 0.00001. (e) Average landmark visibility. The color code corresponds to the percentage of time each landmark was visible at the screen. (f) 3-way interaction effect of Zone, Condition and Landmark on landmark visibility. Each bar shows average landmark visibility (sorted in descending order) for a specific combination of zone (labeled), condition (color) and landmark (texture). We present only combinations associated with at least 10% landmark visibility (17 combinations out of 63). (a, c, e) We divided each trial according to the same sequence of events: walking onset, followed by the first passage in the starting arm (S) then in the finish arm (F), being either the goal or the error arm. Events are horizontally spaced according to the median duration between each event. All three plots represent data in the learning, control, and probe conditions, averaged between separating events across all trials and blocks for all 14 allocentric participants. (b, d, f) Mean value with standard error of the mean (black bars). We present the summary of the significant differences (green braces) found in post-hoc analysis (computed on a pairwise basis, then grouped when similar). For figure (f), we found no pairwise significant differences within the group of combinations not shown (below 10%). *** = p < 0.0001, ** = p < 0.001, * = p < 0.01.

##### Walking speed

Second, we analyzed the walking speed across different conditions and zones (Figs. 3c,d). We found a significant main effect of Zone (*F*(6;273) = 472.15, *p* < 0.00001). Post-hoc analysis revealed that the participants spent more time, and exhibited a slower walking speed at the beginning of the starting arm (i.e., in zone S1, both before and after walking onset, t(2) < −15, *p* < 0.00001 for all pairwise comparisons involving either staticS1 or mobileS1). We observed a tendential, but not significant, main effect of Condition on the walking speed (*F*(2;273) = 3.91, *p* = 0.021, which did not survive the multiple comparisons correction). There was no interaction effect between Condition and Zone (*F*(12;273) = 0.45, *p* = 0.94).

##### Landmark visibility

Third, we tested the visibility of the landmarks depending on the condition, zone and landmark (Fig. 3e) and we observed a 3-way interaction between all factors (*F*(24;819) = 25.31, *p* < 0.00001). Post-hoc analysis (Fig. 3f) revealed a clear tendency for landmarks being visible in the starting arm of the maze (as opposed to the center zone and the finish arm), modulated by the condition and the landmark attended. Figure 3f shows the landmark visibility of [Condition; Zone; Landmark] combinations in descending order, and we can notice steps of combination triplets with the same Zone factor (from staticS1 to C only), in the order in which they are visited by the participants. The consistent pattern in each triplet shows a preferred landmark for each condition: Landmark 1 for learning and control trials, and Landmark 2 for probe trials. A slight deviation from the dominant pattern is that the mean visibility of Landmarks 1 & 2 in Center zone during probe trials are found at the same level (Fig. 3f), although not statistically different from the visibility of any landmark in any condition in the same zone. All additional statistical results (main effects, 2-way interactions) are presented in Supplementary Table 2.

### EEG cluster analysis

#### Independent component selection

To give an overview of the independent component (IC) inspection and selection process, we provide IC and cluster counts at different steps of our procedure (see Fig. 2). In total, we extracted 1943 ICs out of the whole dataset (16 participants). First, at the individual IC labeling step, we relabeled 204 out of 394 ICs initially labeled as ‘Brain’ (i.e., automatic rejection based on RV threshold and manual inspection of hybrid cases). Starting from 1120 input ICs, the clustering algorithm placed 1047 ICs in valid clusters (73 outliers). Then, to complete the cluster labeling step, we selected 35 ‘clusters of interest’ out of the 50 output clusters. We reviewed 755 ICs and edited the label in 207 of them. Eventually, we removed a total of 414 ICs in disagreement with their cluster label, leaving 40±2 ICs per participant (mean ± standard error) for the final analysis (all clusters included).

#### Cluster description

Out of the 50 clusters, we obtained: 2 Eye, 24 Muscle, 12 Brain, 1 Heart, 1 Channel Noise, 0 Line Noise, and 4 Other clusters. The last 6 clusters did not contain a class represented by at least 50% of the ICs.

##### Eye clusters

The 2 Eye clusters contained a mixture of components linked to horizontal and sometimes vertical eye movements. We manually separated these 2 categories, easily identifiable at the IC level, and we focused on the largest ‘Horizontal Eye’ cluster (Fig. 4a). In all conditions, the ERSPs showed a significant increase in horizontal eye movements relative to baseline recordings, during which eyes were supposedly at rest (Fig. 4b). The power increase was particularly pronounced before reaching the Center zone, especially in the lower frequencies (1 Hz to 5 Hz). The significant differences between conditions reflected the increased eye-exploration behavior in the probe condition, mostly before leaving the starting zone S1 (Fig. 4c).

**Figure 4.**
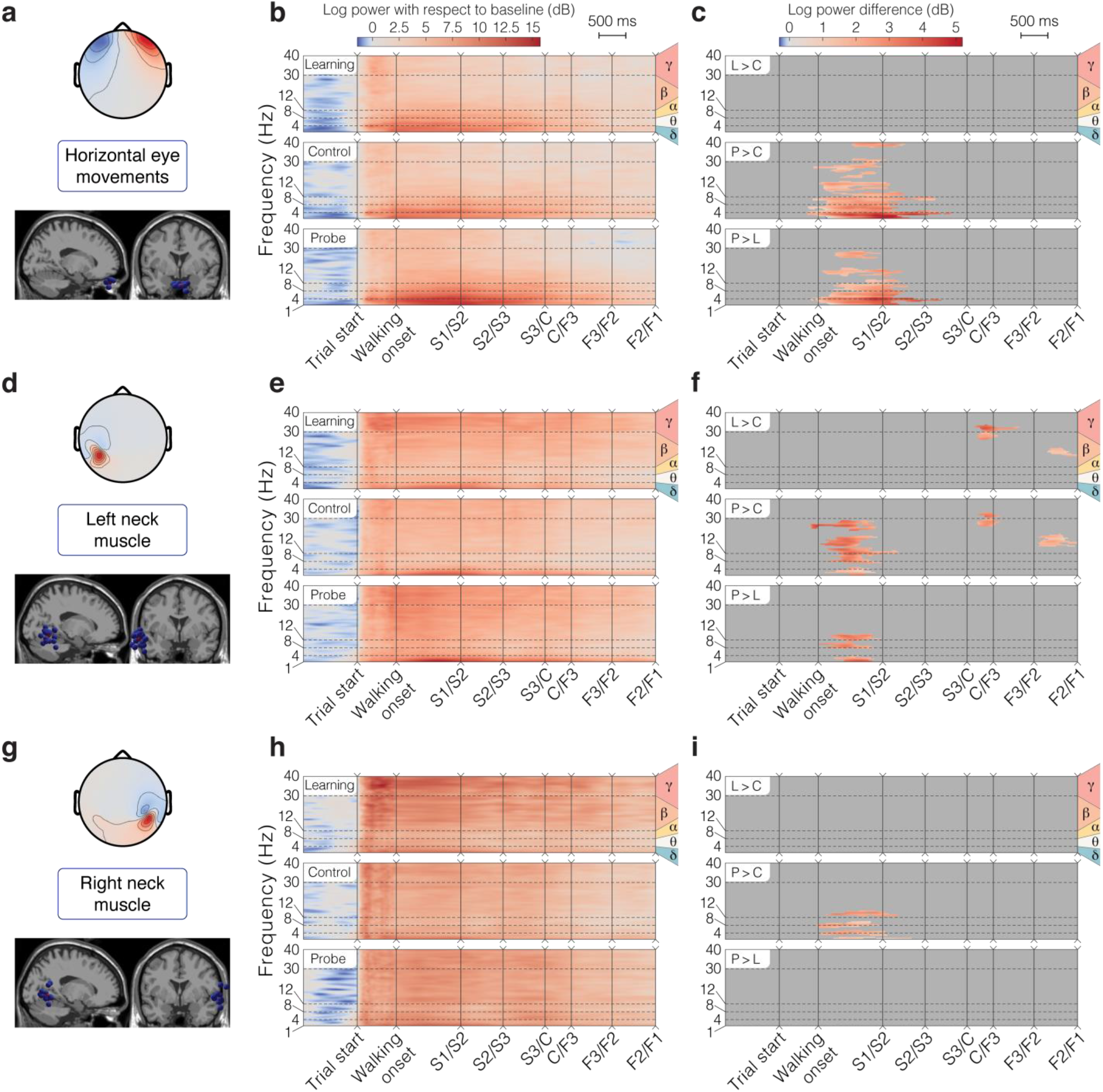
Horizontal eye movements and neck muscle clusters for the allocentric group. **(a, d, g)** Topographical map of the average cluster components’ projection at the scalp level and sagittal view of all ICs in the cluster (blue spheres) with the position of the centroid (red sphere). **(b, e, h)** ERSP average per condition. We first averaged the data at the participant level, then at the group level. **(c, f, i)** ERSP pairwise differences between conditions. Plotted values represent the average of participant-wise ERSP difference between the 2 conditions compared. We masked differences not satisfying the statistical threshold in the permutation test (i.e., p > 0.05). For all the ERSP plots, the Y axis displays the delta, theta, alpha, beta and gamma frequency bands and the X axis represents the time-warped sequence of main events in the trial. We horizontally spaced the events according to the median duration between them. ‘L > C’: difference between Learning and Control, ‘P > C’: difference between Probe and Control, ‘P > L’: difference between Probe and Learning. **(a-c)** Horizontal eye movements cluster. **(d-f)** Left neck muscle cluster. **(g-i)** Right neck muscle cluster.

##### Neck muscle clusters

We identified the neck muscle clusters (out of 24) most likely to reflect sternocleidomastoid activity, based on their topographic activation map and associated dipole location. Two neck muscle clusters (one on each side) were selected (Figs. 4d,g). The ERSPs of both clusters revealed an increased activity of the muscles with respect to the baseline period, across all frequency bands (Figs. 4e,h). The muscle activity in both clusters (high beta and gamma band > 20 Hz; Pion-Tonachini *et al*., 2019) was the greatest after maze appearance and it faded out as the participants walked through the maze. For the left-side cluster, power in mobileS1 was significantly greater in the probe condition than in the other conditions (Fig. 4f), like for the Eye cluster. We also found a significantly increased muscle activity in the learning and probe conditions as compared to control at the center of the maze and just before the end of the task (Fig. 4f). For the right-side cluster, the learning condition seemed to be associated with higher and more sustained activity, but the difference with other conditions was not significant in high frequencies (Figs. 4h,i).

##### Brain clusters

Concerning the brain clusters, we kept only those containing ICs coming from at least 9 of the 14 allocentric participants (~65%), to ensure that they were representative enough of our sample population (detailed information about the 12 brain clusters is reported in Supp. Table 4). This sorting left 6 clusters for analysis. Using the Talairach client (Lancaster *et al*., 2000), we computed the closest gray matter region to each brain cluster centroid. As shown in Figure 5, the 6 selected clusters of interest were located in or near BA23 in the posterior cingulate (Cluster 1: [8,-47,25]), BA19 in the right cuneus overlapping with BA7 in the right precuneus (Cluster 2: [15,-82,35]), BA40 in the right supramarginal gyrus (Cluster 3: [39,-51,33]), BA33 in the anterior cingulate (Cluster 4: [-2,9,22]), BA6 in the right precentral gyrus (Cluster 5: [33,-9,52]), and BA3 in the left postcentral gyrus (Cluster 6: [-37, −28,49]). These coordinates ([x,y,z]) are in Talairach units.

**Figure 5.**
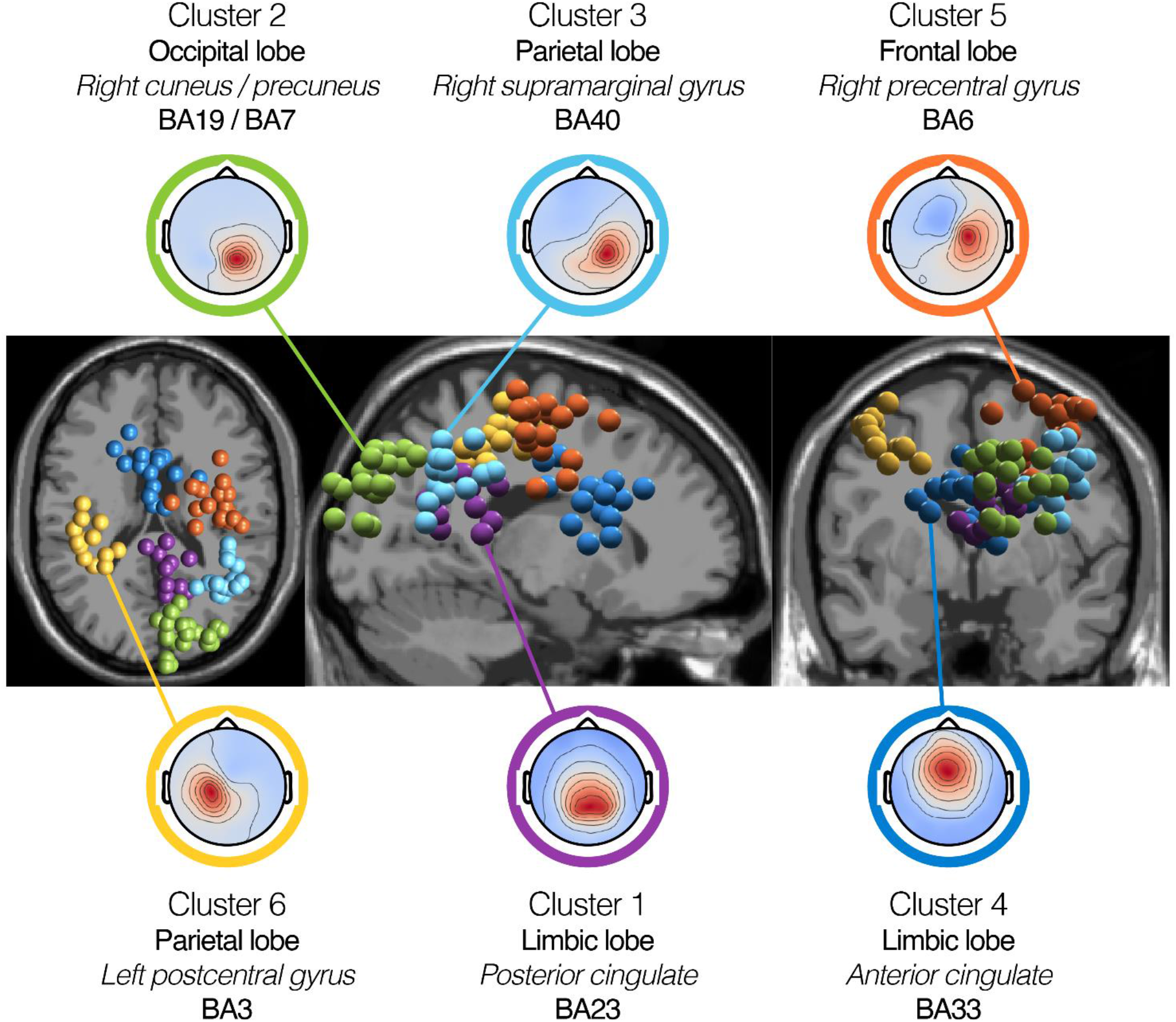
Brain cluster 3D localization and mean channel activation maps. Spatial location of brain clusters retained for analysis (from left to right: transverse view, sagittal view, coronal view). Each IC is represented by a sphere located at its corresponding dipole location. For each cluster, we plotted all ICs, irrespective of their associated participant’s behavioral group. We used MRI scans from the standard MNI brain for representation. Topographies show the mean channel activation map associated to each cluster. The centroids of the clusters are located in or near the posterior cingulate (Cluster 1 – 12 ICs, 12 participants), the right cuneus (Cluster 2 – 22 ICs, 12 participants), the right supramarginal gyrus (Cluster 3 – 15 ICs, 11 participants), the anterior cingulate (Cluster 4 – 15 ICs, 12 participants), the right precentral gyrus (Cluster 5 – 17 ICs, 13 participants), the left postcentral gyrus (Cluster 6 – 13 ICs, 11 participants). Detailed information on the location of the cluster centroids is provided in Supplementary Table 4.

#### Brain cluster activity

The analyses of the 6 selected brain clusters are presented in Figures 6 and 7 (clusters 1-3 and 4-6, respectively).

**Figure 6.**
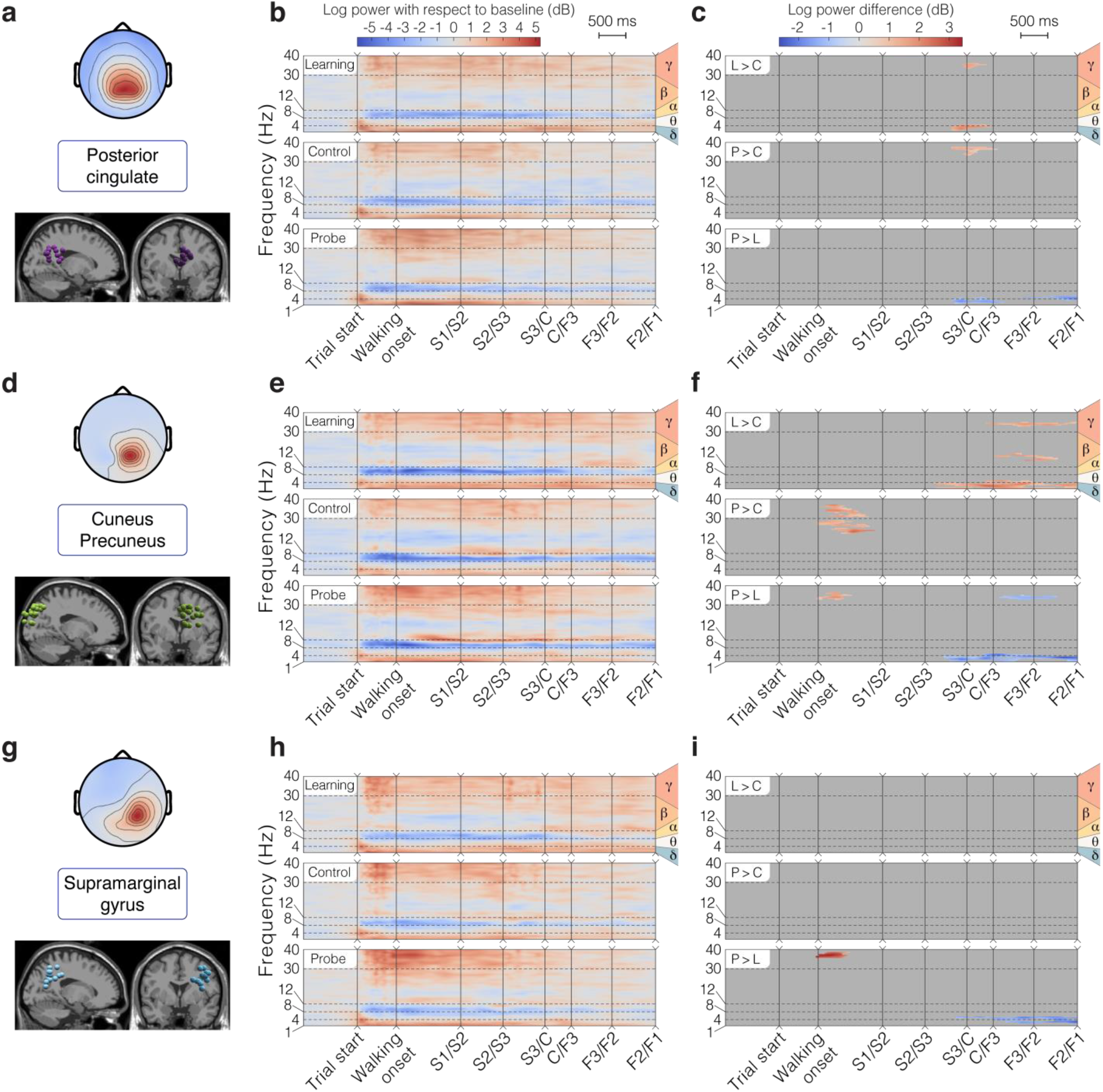
Detailed analysis of brain clusters 1 – 3 for the allocentric group. **(a, d, g)** Topographical map of the average cluster components’ projection at the scalp level (top) and sagittal/frontal views of all ICs in the cluster (bottom). **(b, e, h)** ERSP average per condition. **(c, f, i)** ERSP pairwise differences between conditions. ‘L > C’: difference between Learning and Control, ‘P > C’: difference between Probe and Control, ‘P > L’: difference between Probe and Learning. **(a-c)** Cluster 1 – Posterior Cingulate. In the allocentric group, this cluster contains 10 ICs from 10 different participants. **(d-f)** Cluster 2 – Right Cuneus/Precuneus. In the allocentric group, this cluster contains 21 ICs from 11 different participants. **(g-i)** Cluster 3 – Right Supramarginal Gyrus. In the allocentric group, this cluster contains 13 ICs from 9 different participants.

**Figure 7.**
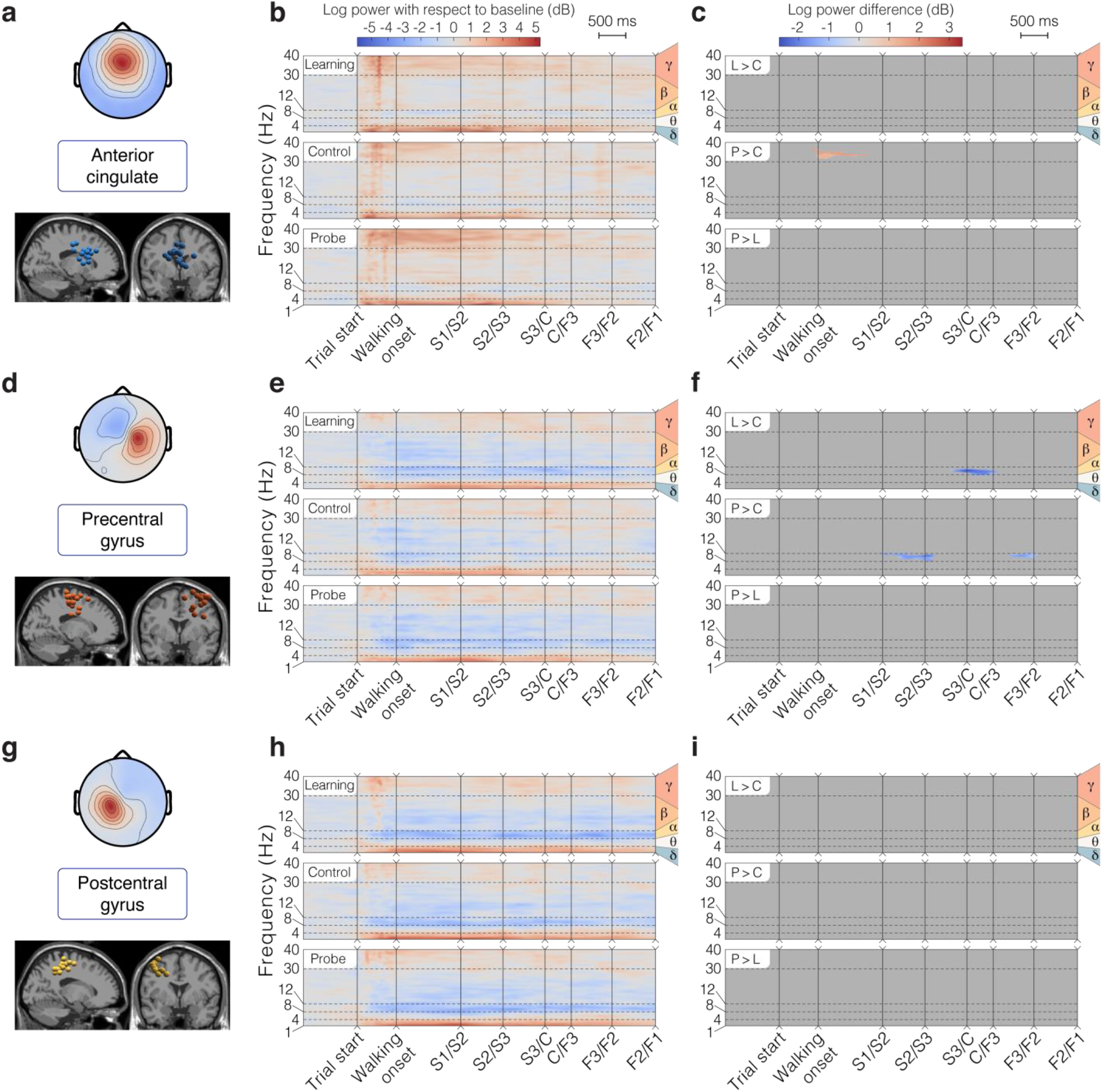
Detailed analysis of brain clusters 4 - 6 for the allocentric group. The layout is the same as in Fig. 6. **(a, d, g)** Topographical map of the average cluster components’ projection at the scalp level (top) and sagittal/frontal views of all ICs in the cluster (bottom). **(b, e, h)** ERSP average per condition. **(c, f, i)** ERSP pairwise differences between conditions. ‘L > C’: difference between Learning and Control, ‘P > C’: difference between Probe and Control, ‘P > L’: difference between Probe and Learning. **(a-c)** Cluster 4 – Anterior Cingulate. In the allocentric group, this cluster contains 14 ICs from 11 different participants. **(d-f)** Cluster 5 – Right Precentral Gyrus. In the allocentric group, this cluster contains 15 ICs from 11 different participants. **(g-i)** Cluster 6 – Left Postcentral Gyrus. In the allocentric group, this cluster contains 12 ICs from 10 different participants.

##### Alpha band activity (8-12 Hz)

The average ERSP analysis for posterior parieto-occipital clusters (1-3) showed a marked alpha (8-12 Hz) desynchronization (power suppression of 3 dB or more) starting after trial onset in all conditions (Figs. 6b,e,h). This power suppression slowly faded away or narrowed down around 9 Hz when the participant left the first section of the maze. The desynchronization was less marked in the precentral and the postcentral gyri, but it was sustained throughout the trial, except for the control condition (significant difference found for the precentral gyrus near the central zone of the maze, see Fig. 7f). In the anterior cingulate, we intermittently observed a similar but reduced alpha power suppression (difference of 1 dB with respect to baseline, Fig. 7b).

##### Gamma band activity (>30 Hz)

We found that gamma (> 30Hz) synchronization was strongly enhanced in this navigation task, with clusters 1-4 (posterior and anterior cingulate, cuneus, supramarginal gyrus) presenting amplitudes greater than baseline in this frequency band, consistently throughout the maze (Figs. 6,7). Nonetheless, a power increase in this frequency band was found between trial start and the center, especially in the probe condition (significant differences found in the mobileS1 zone for cuneus, supramarginal gyrus and anterior cingulate clusters). A comparison between conditions also demonstrated a reduced gamma activity upon reaching the center in the control condition in the posterior cingulate and an increased gamma power in the learning condition in the cuneus in the finish arm.

##### Delta and theta band activity (<8 Hz)

Finally, we observed modulations of low frequency rhythms (delta range 1-4 Hz, theta range 4-8 Hz), with sustained greater delta amplitudes in the starting arm in all brain clusters and a strong transient theta burst at the beginning of the trial in posterior parieto-occipital clusters (1-3). The brain activity in these frequency bands proved to be condition specific for these clusters, with a generally higher power for the learning condition along the finish arm (Figs. 6c,f,i).

## Discussion

This work brings together the technology and data analysis tools to perform simultaneous brain/body imaging during landmark-based navigation in fully mobile participants. Our behavioral results show that a majority of young adults can rapidly learn to solve the Y-maze by using an allocentric strategy, confirming previous findings in similar landmark-based navigation studies (Kimura *et al*., 2019; Bécu *et al*., 2020b). We find that allocentric participants have the capacity to flexibly reorient by observing landmarks at the beginning of the trial (consistent with the more precise gaze dynamics described in Bécu *et al*., 2020b). The analysis of high-density EEG data shows that exploitable neural signals are extracted from various brain regions (posterior cingulate, cuneus/precuneus, supramarginal gyrus, anterior cingulate, precentral gyrus, and postcentral gyrus) that replicate and extend previous neuroimaging findings from a similar fMRI study (Ramanoël, Durteste, *et al*., 2020). Overall, the identified brain structures represent an extended ensemble of areas involved in the high-level processing of visual information, in spatial representation, and in motor planning necessary to navigate.

### Task-solving behavior

The zone-based analysis reveals a common task-solving behavior in allocentric participants, starting by an observation period at the beginning of the trial (slow speed, high variability in horizontal heading, and maximal visibility of landmarks) followed by navigation to the chosen arm. We found minimal head rotations after reaching the center of the maze and no significant deceleration, indicating the initial observation period to be the main source of visual information for the participants. This interpretation was also supported by the analysis of horizontal eye movement and neck muscle activity clusters, which exhibited greater activity at the beginning of the maze (Figs. 4b,e,h). Interestingly, between-condition contrasts revealed an accentuation of this pattern in the probe condition (Figs. 4c,f), probably reflecting a higher need for information gathering at an unfamiliar starting point.

The landmark visibility analysis confirmed that navigators may rely extensively on the landmark appearing straight ahead at the beginning of the task, which differs in the probe condition (Landmark 2) from the other conditions (Landmark 1). This suggests that the participants were mainly capable of reorienting with the information gathered from this landmark only, even in the probe condition. We found additional tendencies for this metric (see last columns of Fig. 3f), such as (i) a similar visibility for the two predominant landmarks at the center during probe condition and (ii) an increased visibility for the second most visible landmark (relative to each condition) with respect to the third during the initial observation period in the learning and probe conditions. None of these observations were applicable to the two egocentric participants (Supp. Fig. 2e,f). Therefore, these findings might reflect a perceptual mechanism helping to bind multiple landmarks in a single representation of the environment, specific to the allocentric participants’ strategy.

### Anatomical substrates of the clusters retrieved

We hypothesized that the analysis of brain dynamics in fully mobile individuals would retrieve structures involved in active, multimodal landmark-based spatial navigation. Here, we contrast our results against those from static neuroimaging paradigms. First, we expected the retrosplenial complex (RSC) to play a central role in solving the Y-maze, as this task requires landmark-based re-orientation. Accordingly, we found a cluster located in or near the posterior cingulate cortex (cluster 1), encompassed by the RSC (Julian *et al*., 2018). fMRI studies have consistently shown that the human RSC encodes heading direction (Marchette *et al*., 2014; Shine *et al*., 2016) anchored to local visual cues, like stable landmarks, by using a first-person perspective (Auger *et al*., 2012; Marchette *et al*., 2015; Auger & Maguire, 2018). Moreover, the RSC is embedded in a network of somatosensory areas that were also partially retrieved in our cluster analysis. In particular, the precuneus (cluster 2) is involved in several aspects of spatial cognition, such spatial attention, spatial working, long-term memory, and the representation of landmarks, in association with RSC during navigation (Cona & Scarpazza, 2019). Also, in line with the mobile aspects of our paradigm, several studies have highlighted the role of the precuneus in the integration and coordination of motor behavior during navigation (Navarro *et al*., 2018). Around the centroid of the cluster associated to the precuneus, we note that the independent components (ICs) forming the cluster are distributed across the anatomical boundaries between occipital and parietal cortices. This region spans areas mediating visuo-spatial processing, such as the Occipital Place Area, which is sensitive to navigable pathways in a perceived scene (Bonner & Epstein, 2017; Patai & Spiers, 2017).

Our EEG analysis also retrieved three clusters associated to the supramarginal gyrus (cluster 3), the anterior cingulate cortex (cluster 4), and the precentral gyrus (cluster 5). The supramarginal gyrus, which belongs to the somatosensory cortex, plays a role in the mnemonic components of spatial navigation (van der Linden *et al*., 2017; Sneider *et al*., 2018). In addition, as it is encompassed within the inferior parietal lobule, the supramarginal gyrus is involved in spatial attention (Cona & Scarpazza, 2019). The anterior cingulate cortex subserves high-level cognitive functions such as route planning (Spiers & Maguire, 2006; Lin *et al*., 2015) as well as its re-evaluation and updating based on internal monitoring, more specifically with respect to error detection and spatial reorientation (Javadi *et al*., 2019). As for the right precentral gyrus cluster, we report its centroid in BA6 although the spatial extent of its ICs spans towards more frontal areas. Considering this limited spatial precision, we speculate a putative contribution from the supplementary motor area proper and the right middle frontal gyrus, both overlapping with BA6. The former is recruited in motor planning (Simon *et al*., 2002) and motor execution of self-initiated movements (Cona & Semenza, 2017), consistent with the mobile aspect of our task. The latter is involved in spatial attention and spatial working memory, specifically in BA6 (Cona & Scarpazza, 2019). Finally, our brain analysis retrieved postcentral gyrus activity (cluster 6), encompassing the primary somatosensory cortex, and thus most likely to be involved in the processing of proprioception (Rausch *et al*., 1998).

### Functional analysis of the clusters’ activity

#### Gamma band activity (>30 Hz)

An objective of this work was to couple brain and body imaging during spatial navigation. Complementing the behavioral findings, the analysis of transient time-frequency EEG patterns shows a strong gamma band synchronization in posterior parieto-occipital clusters, especially in the starting arm (Figs. 6b,e,h), which coincides with the increased eye movement related activity observed in the same spatial area (low-frequencies in Fig. 4b). In line with findings showing that increased gamma power in parieto-occipital region promotes sharper visuo-spatial attention (Gruber *et al*., 1999; Müller *et al*., 2000; Jensen *et al*., 2007), this cortical activity pattern supports our interpretation of the participants’ behavior. We reported significant differences when the participant starts walking in the probe condition compared to the learning and control conditions (in the cuneus/precuneus, Fig. 6f, and supramarginal gyrus, Fig. 6i). This pattern may reflect a greater attentional demand triggered by the visual conflict between the probe and the other conditions, forcing the participant to actively reorient. Statistical analyses conducted on eye and muscle clusters also revealed a more active state (more frequent eye movements and increased muscle activity) when the participant starts walking during probe trials. The greater involvement of posterior parietal cortex (especially the precuneus) during this crucial reorientation moment is coherent with the fMRI evidence linking it to the navigationally-relevant representation of landmarks, when participants are moving with respect to stable objects (Cona & Scarpazza, 2019). The mobile EEG literature of locomotion control more often reports activity bound to steady-state gait cycle events (e.g., Gwin *et al*., 2011; Castermans *et al*., 2014; Wagner *et al*., 2014, 2016; Luu *et al*., 2017a,b), making it difficult to compare with our experimental design. Several works presenting results contrasting a walking condition with a standing baseline condition described a desynchronization in the high beta band (25-35 Hz) in the sensorimotor cortex (Wagner *et al*., 2012; Seeber *et al*., 2014, 2015), which does not concur with our findings (Figs. 7e,h). Nonetheless, Bulea *et al*. (2015) report high gamma band synchronization (30-50 Hz) when comparing active walking to quiet standing in the posterior parietal area, which better aligns with our results (Figs. 6b,e,h). However, locomotor control can only be a part of the interpretation since it does not explain the specific activity observed in the probe condition.

#### Alpha band activity (8-12 Hz)

Our ERSP analysis shows a desynchronization in the alpha band, spanning almost all clusters (except the anterior cingulate), with different temporal dynamics. In the sensorimotor cortex (post-central and pre-central gyri), the desynchronization extends to the low beta band, it starts a few moments before movement onset and it is sustained throughout the whole maze traversal (Figs. 7e,h). This pattern advocates for a mere signature of locomotion, as reported in numerous mobile EEG studies comparing walking and standing (Presacco *et al*., 2011; Seeber *et al*., 2014; Bulea *et al*., 2015). Further supporting the idea that such activity is a neural correlate of ambulation, no differences between conditions were found in postcentral gyrus (Fig. 7i). Interestingly, the precentral gyrus exhibited a modulation of activity around the center of the maze where alpha desynchronization was less pronounced in the control condition as compared to others (Fig. 7f). The precentral gyrus is known to be associated with movement planning (Wagner *et al*., 2014; Navarro *et al*., 2018). Alpha power suppression has been linked to increased activity in motor regions (Pfurtscheller & Klimesch, 1991), such that this activity pattern could reflect a more passive execution of the turn in a situation in which the participant can straightforwardly repeat the learning condition. Although this purely ambulatory feature extends to more posterior parietal areas (Bulea *et al*., 2015), the temporal dynamics of the alpha power in our parieto-occipital clusters (i.e., an almost immediate desynchronization after trial start and a subsequent fading across maze traversal, Figs. 6b,e,h) suggests a different interpretation. According to the meta-analysis from Cona & Scarpazza (2019), the precuneus and the inferior parietal lobule are embedded in a fronto-parietal network mediating spatial attention. Thus, the fading of the desynchronization might reflect a progressive decrease in spatial attention, as sufficient visual information is being gathered. As participants seem to make their decision early in the task, they should reach their maximal degree of alertness in the first sections of the maze and let it drop afterwards. Echoing this interpretation, several EEG studies of spatial navigation associated alpha power in the parietal cortex to spatial learning (Gramann *et al*., 2010; Lin *et al*., 2015), with significant task-related modulations.

In an experiment reporting the modulation of RSC activity in passive simulated navigation, those participants who relied on an allocentric reference frame demonstrated a sustained alpha power decrease during straight segments and a strong alpha power increase during absolute heading rotation (Chiu *et al*., 2012; Lin *et al*., 2015, 2018). For our posterior cingulate cluster, such heading discriminant activity is neither observed at the starting position where head movements are maximal (relative to the body) nor near the central zone of the maze (relative to the global environment, i.e., the landmarks) (Fig. 6b). Partially explaining these diverging results, Gomez *et al*. (2014) reported a stronger RSC activation during on-the-spot rotation as compared to continuous movement, tempering the heading computation role of RSC when translational movements are involved. Additionally, the alpha desynchronization elicited by a desktop-based rotation is absent when performed physically (Gramann *et al*., 2018), which shows the important influence of vestibular and proprioceptive cues in modulating RSC activity. Thus, assuming that our posterior cingulate cluster is bound to RSC activity, our results provide additional evidence that the involvement of RSC in heading calculation has been overestimated with respect to ecological navigation. The dynamics of the posterior cingulate cluster in the alpha frequency band are more in accordance with a memory role serving the encoding/retrieval of the egocentric percepts into the allocentric representation (Vann *et al*., 2009; Mitchell *et al*., 2018). Indeed, the fact that alpha desynchronization occurred during the observation of the environment suggests the association of RSC with the encoding/decoding of landmark-based information. This interpretation fits with the tendency of our cluster to be localized in the dorsal part of the posterior cingulate, which is known to play a role in spatial recall tasks in opposition to the ventrolateral part, more likely to be activated during tasks proposing passive viewing or active navigation without the need to respond, perform spatial computation, or self-localize (Burles *et al*., 2018).

#### Delta and theta band activity (<8 Hz)

We observe a strong delta band synchronization (< 5 Hz) in all clusters, lasting the whole traversal of the maze, which has been previously reported as a motion-related artifact (Gwin *et al*., 2010; Presacco *et al*., 2011; Castermans *et al*., 2014). However, the presence of condition-specific modulations in posterior parieto-occipital clusters casts a doubt on this interpretation. During the learning condition, statistical analyses demonstrated a sustained delta/theta synchronization in the finish arm (starting in the center zone, Figs. 6c,f,i). Possibly elucidating this feature, a previous study of spatial working memory in mobile conditions observed a similar theta synchronization seconds prior to the stimulus presentation in posterior cingulate and somatosensory association areas (Kline *et al*., 2014). Arguing that theta power modulations can be related to memory encoding and maintenance, this may be the signature of a learning mechanism, preparing to encode the outcome of the learning trial at the end of the finish arm. However, unlike Kline *et al*. (2014), we do not find subsequent theta desynchronization on stimulus presentation (goal appearance). Another deviation from the artifactual hypothesis is that the delta/theta synchronization seems specific to the starting arm for the anterior cingulate cortex (Fig. 7b), coinciding with participants’ decision-making period as indicated by behavioral analyses. This may reflect the increased spatial working memory demand required for route planning, since previous studies reported increased theta power in the frontal cortex during more cognitively demanding navigation periods (Kahana *et al*., 1999; Caplan *et al*., 2003; Lin *et al*., 2015). Closer to the interrogations posed by the present task, Ferguson *et al*. (2019) found the anterior cingulate to mediate a reinforcement learning role by eliciting a reward when allocentric navigators were shown previously learned cues predicting the goal location. In addition, we observed theta bursts of activity (4-8 Hz) closely time-locked to the beginning of the task in most clusters (mainly in the posterior cingulate, Fig. 6b, and supramarginal gyrus, Fig. 6h). This pattern of activity may be framed within a postural control interpretation: as the environment suddenly appears to the participant, his/her balance control system, previously deprived of any visual information, needs to be updated based on the novel visual cues (Flückiger & Baumberger, 1988; Horak & Macpherson, 2011). Strikingly, theta bursts of activity were similarly described immediately following spontaneous loss of balance from walking on a beam (Sipp *et al*., 2013) and sudden visual perturbations to standing or walking balance (Peterson & Ferris, 2018). These bursts were noticeable in posterior cingulate and posterior parietal areas, associated with vestibular sensing (Kim *et al*., 2017) and resolving visual conflicts (Peterson & Ferris, 2018), respectively.

### Limitations

Source reconstruction was performed using an electrodes’ location template and average MRI anatomical data, which limited its spatial accuracy. Thus, the interpretations proposed in this work should be treated with caution. The use of subject-specific data to build the head model would help increase the accuracy of source localization algorithms (Akalin Acar & Makeig, 2013; Shirazi & Huang, 2019) and it would eventually enable more robust interpretations of the neural correlates of spatial behavior.

Although the methods employed here to clean the EEG signals have been previously validated in the literature (Nordin *et al*., 2019; Richer *et al*., 2019), there is never complete guarantee that the results are artifact free. In particular, the muscular activity associated with microsaccades has been shown to resist standard cleaning methods (Yuval-Greenberg *et al*., 2008; Hassler *et al*., 2011) and it could in principle be contributing to the brain ICs in the gamma frequency band (Yuval-Greenberg *et al*., 2008). However, the influence of this type of artifact is meaningful in experimental setups favoring the accumulation of microsaccades at a fixed latency with respect to the synchronizing event (e.g., fixation of a visual target; Yuval-Greenberg *et al*., 2008) and the probability that this applies to our setup is low.

Gait-related artifact contamination is another well-known pitfall of ambulatory studies (Castermans *et al*., 2014). Walking induces small motions of electrodes and cables that can have a large impact on the signal-to-noise ratio. Typically, the spectral signature of such artifacts contains elevated power amplitudes at the stepping frequency (between 0.5 and 1 Hz for normal walking speeds) and its harmonics, as well as a power modulation pattern time-locked to gait cycle especially marked below 20 Hz (Castermans *et al*., 2014; Snyder *et al*., 2015). Therefore, as already acknowledged, the low frequencies (between 1 and 5 Hz) power increase observed consistently through our clusters may reflect this type of contamination. Yet, motion artifacts were found negligible during treadmill walking at moderate speeds such as those adopted by participants in our study (Gwin *et al*., 2010; Nathan & Contreras-Vidal, 2016). Also, the wireless property of our EEG system (as in Nathan & Contreras-Vidal, 2016) provides additional robustness to gait-related artifacts by minimizing cable sways, identified as major artifactual causes (Symeonidou *et al*., 2018).

Considering the recent publication of the ICLabel algorithm, our work provides some practical insights on how to integrate this promising tool into the MoBI approach. At first, the automatic IC classification has proven to be very useful to deal with large numbers of components. However, through the manual inspection of a large proportion of the automatically assigned labels, we uncovered and corrected a substantial amount of discrepancies between the algorithm and the experimenter’s opinion, thus adopting a semi-automated procedure. Even if human categorization of ICs can be variable and error prone (Pion-Tonachini *et al*., 2019), we believe that these discrepancies also stem from complex artifact patterns present in mobile EEG. However, resorting to the experimenter’s judgement is not desirable for future studies since it impairs replicability and it is very time-consuming for high-density recordings. Future works should explore the flexibility of interpretation offered by the compositional label, for example by adapting the probability thresholds to better tailor the algorithm’s output to the characteristics of the data being processed.

## Conclusion & Future works

This study provides a proof-of-concept about the possibility of imaging the neural bases of landmark-based spatial navigation in mobile, ecological set-ups. First, the presented EEG analysis identifies a set of brain structures also found in fMRI studies of landmark-based spatial cognition. Second, our approach reveals the role of brain areas involved in active, fully engaging spatial behavior (such as clusters in the sensorimotor cortex related to motor execution and proprioception), whose contribution is usually overlooked in static fMRI paradigms. We present new insights onto the cortical activity mediating successful spatial reorientation when visual, proprioceptive and vestibular sensory inputs are coherent. Specifically, alpha band desynchronization in the posterior cingulate when participants gather visual information provides further support to the idea that RSC plays an important role at the interface between perception of landmarks and spatial representation. Despite showing few effects of experimental condition, our results illustrate the benefit, in terms of deciphering neural dynamics within the course of a trial, of fine temporal resolution brain imaging paired with meaningful behavioral markers during spatial navigation.

The methodology associated to the MoBI approach remains quite new and such experiments help to identify vectors of improvement. At the preprocessing stage, further characterization of the parameters and robustness comparison with other pipelines (such as Automagic, Pedroni *et al*., 2019) would be beneficial. Complementary steps such as sliding window approaches for isolating transient artifacts using principal component analysis and/or canonical correlation analysis can improve source separation compared to ICA alone (Artoni *et al*., 2017; Nordin *et al*., 2020). Adding simultaneous noise and neck electromyographic recordings have also been shown to successfully assist the identification and removal of motion-related artifacts (Nordin *et al*., 2019, 2020). Concerning the gathering of insights on strategy-specific behaviors, additional improvements of the protocol are also desirable. Using a passively guided traversal of the maze as a baseline to contrast with the main task may help to disentangle the neural correlates of locomotion control and active landmark-based spatial navigation. The addition of an eye tracking system embedded in the VR head-mounted display would also bring further insights on the differential role of visuo-spatial cues (Bécu *et al*., 2020a).

## Acknowledgments

We first thank the participants of this study for their patience and precious contribution. We thank Elisa Tartaglia for the helpful discussions along the project, and Benjamin Paulisch for his great help in setting up the experiment. We also deeply thank Marion Durteste for her text proof-reading. This research was supported by the Chair SILVERSIGHT ANR-14-CHIN-0001 & ANR-18-CHIN-0002, the LabEx LIFESENSES (ANR-10-LABX-65), and the IHU FOReSIGHT ANR-18-IAHU-01.

## Competing Interests

The authors declare no conflict of interest.

## Author Contributions

AD, JBDSA, SR, MB, RC, KG, and AA designed the experiment. AD and JBDSA conducted the experiment and collected the data. AD, JBDSA, LG, and MK analyzed the data. AD, JBDSA, SR, and AA wrote the manuscript. AD, JBDSA, SR, MB, LG, MK, RC, JAS, KG, and AA revised the manuscript.

## Data Accessibility Statement

Data and scripts used for the analysis are available by request to the corresponding authors (Alexandre Delaux; Jean-Baptiste de Saint Aubert).

## Abbreviations

AMICA: Adaptative Mixture Independent Component Analysis;
ANOVA: Analysis of Variance;
BA: Brodmann Area;
ECD: Equivalent Current Dipole;
EEG: Electroencephalogram;
ERSP: Event-Related Spectral Perturbations;
fMRI: functional Magnetic Resonance Imaging;
IC: Independent Component;
ICA: Independent Component Analysis;
MNI: Montreal Neurological Institute;
MoBI: Mobile Brain/Body Imaging;
ROI: Region of Interest;
RSC: Retrosplenial Complex;
VR: Virtual Reality;

## Footnotes

1 In 3D virtual reality and computer graphics, the viewing frustum is defined as the region of virtual space displayed on the screen, and it is a coarse imitation of the “cone of vision” in natural viewing. It takes the form of a truncated rectangular pyramid, defined by the horizontal and vertical field of view and by near and far bounds.

## Notes

<u>Funding:</u> This research was supported by the Chair SILVERSIGHT ANR-14-CHIN-0001 & ANR-18-CHIN-0002, the LabEx LIFESENSES (ANR-10-LABX-65), and the IHU FOReSIGHT ANR-18-IAHU-01.

### Competing Interest Statement

The authors have declared no competing interest.

### Summary of Updates

Introduction updated to state goals and hypotheses more clearly; Flowchart added as Figure 2; Previous Figure 2 moved to Supplementary materials; Removal of unexploited ERP results; Behavioral results discussion added; authors affiliations updated.

