## Supplementary Material for "Mobile brain/body imaging of landmark-based navigation with high-density EEG"

Manuscript proposed for the EJN special issue:
**Time to move: brain dynamics underlying natural action and cognition**

### Mobile brain/body imaging of landmark-based navigation with high-density EEG

#### Supplementary information

Alexandre Delaux[*1], Jean-Baptiste de Saint Aubert[*1], Stephen Ramanoël[1], Marcia Bécu[1], Lukas Gehrke[2], Marius Klug[2], Ricardo Chavarriaga[3,4], José-Alain Sahel[1,5,6,7], Klaus Gramann[2], Angelo Arleo[1]

[*]These authors contributed equally (co-first authorship)

[1]Sorbonne Universités, INSERM, CNRS, Institut de la Vision, Paris, France

[2]Institute of Psychology and Ergonomics, Technische Universität Berlin, Germany

[3]Center for Neuroprosthetics, Ecole Polytechnique Fédérale de Lausanne, Geneva, Switzerland

[4]Zurich University of Applied Sciences, ZHAW Datalab, Winterthur, Switzerland

[5]CHNO des Quinze-Vingts, INSERM-DGOS CIC 1423, Paris, France

[6]Fondation Ophtalmologique Rothschild, Paris, France

[7]Department of Ophthalmology, The University of Pittsburgh School of Medicine, Pittsburgh, United States

***Keywords:*** *mobile EEG, virtual reality, ecological navigation, source reconstruction, retrosplenial complex (RSC)*

This document contains 21 pages, including 4 Methods sections, 4 Figures, and 4 Tables.

#### Supplementary Table 1. *Repartition of trials kept for analysis*

| Participant | Behavioral group | Total number of trials | Learning Trials | Control Trials | Probe Trials |
|---|---|---|---|---|---|
| **P01** | Allocentric | 86 (91) | 33 (37) | 26 (27) | 27 (27) |
| **P02** | Allocentric | 85 (89) | 32 (35) | 27 (27) | 26 (27) |
| **P03** | Allocentric | 88 (90) | 34 (36) | 27 (27) | 27 (27) |
| **P04** | Allocentric | 80 (88) | 27 (34) | 26 (27) | 27 (27) |
| **P05** | Egocentric | 80 (84) | 26 (30) | 27 (27) | 27 (27) |
| **P06** | Egocentric | 80 (84) | 27 (30) | 27 (27) | 26 (27) |
| **P07** | Allocentric | 73 (86) | 25 (32) | 27 (27) | 21 (27) |
| **P08** | Allocentric | 81 (84) | 27 (30) | 27 (27) | 27 (27) |
| **P09** | Allocentric | 79 (88) | 26 (34) | 26 (27) | 27 (27) |
| **P10** | Allocentric | 81 (89) | 27 (35) | 27 (27) | 27 (27) |
| **P11** | Allocentric | 78 (84) | 25 (30) | 27 (27) | 26 (27) |
| **P12** | Allocentric | 81 (89) | 28 (35) | 27 (27) | 26 (27) |
| **P13** | Allocentric | 82 (90) | 28 (36) | 27 (27) | 27 (27) |
| **P14** | Allocentric | 79 (88) | 27 (34) | 27 (27) | 25 (27) |
| **P15** | Allocentric | 76 (85) | 22 (31) | 27 (27) | 27 (27) |
| **P16** | Allocentric | 80 (85) | 26 (31) | 27 (27) | 27 (27) |
| Total Allocentric group | | 1129 (1226) | 387 (470) | 375 (378) | 367 (378) |
| Global dataset | | 1289 (1394) | 440 (530) | 429 (432) | 420 (432) |

***Repartition of the trials per participant kept for the zone-based and EEG analyses.*** We indicate in parentheses the initial number of trials before rejection. We discarded outlier trials that did not comply with the chosen sequence of events and those lasting too long to be consistently incorporated in the analysis. The effective duration cut-off (computed from the distribution of escape latency) was 12956 ms.

### Supplementary Figure 1. *Group behavioral results*

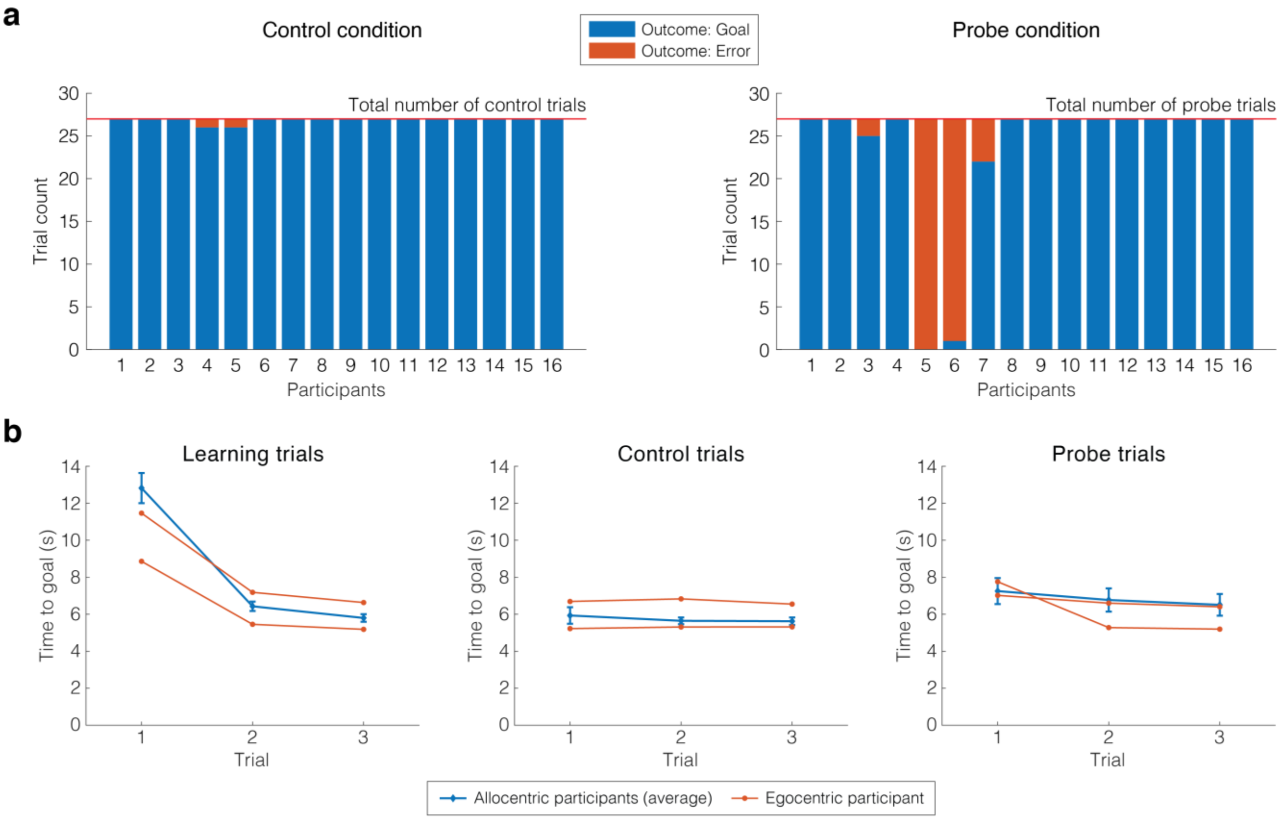

***Behavioral results – Group behavior across trials.* (a)** *Strategy assignment.* Outcome of the control (left) and probe (right) trials showing the participant-wise count of Goal and Error arm choices. Each test part of the experiment (control and probe conditions) comprised a total of 27 trials across blocks. Probe trial outcomes were used to assign each participant a strategy preference: 14 participants had a majority of allocentric responses (choosing the Goal arm in probe trials) and 2 participants had a majority of egocentric responses (choosing the Error arm in probe trials). **(b)** *Group-level time to goal per condition.* The evolution of the time to goal across trials, presented for each condition (learning, control, probe) and averaged across blocks. Data were averaged across participants in the allocentric group (bars indicate standard error of the mean). For the two egocentric participants, individual data are showed. For the learning trials, we considered the first 3 trials, irrespectively of their outcome.

#### **Supplementary Figure 2.** ***Egocentric participants' behavior***

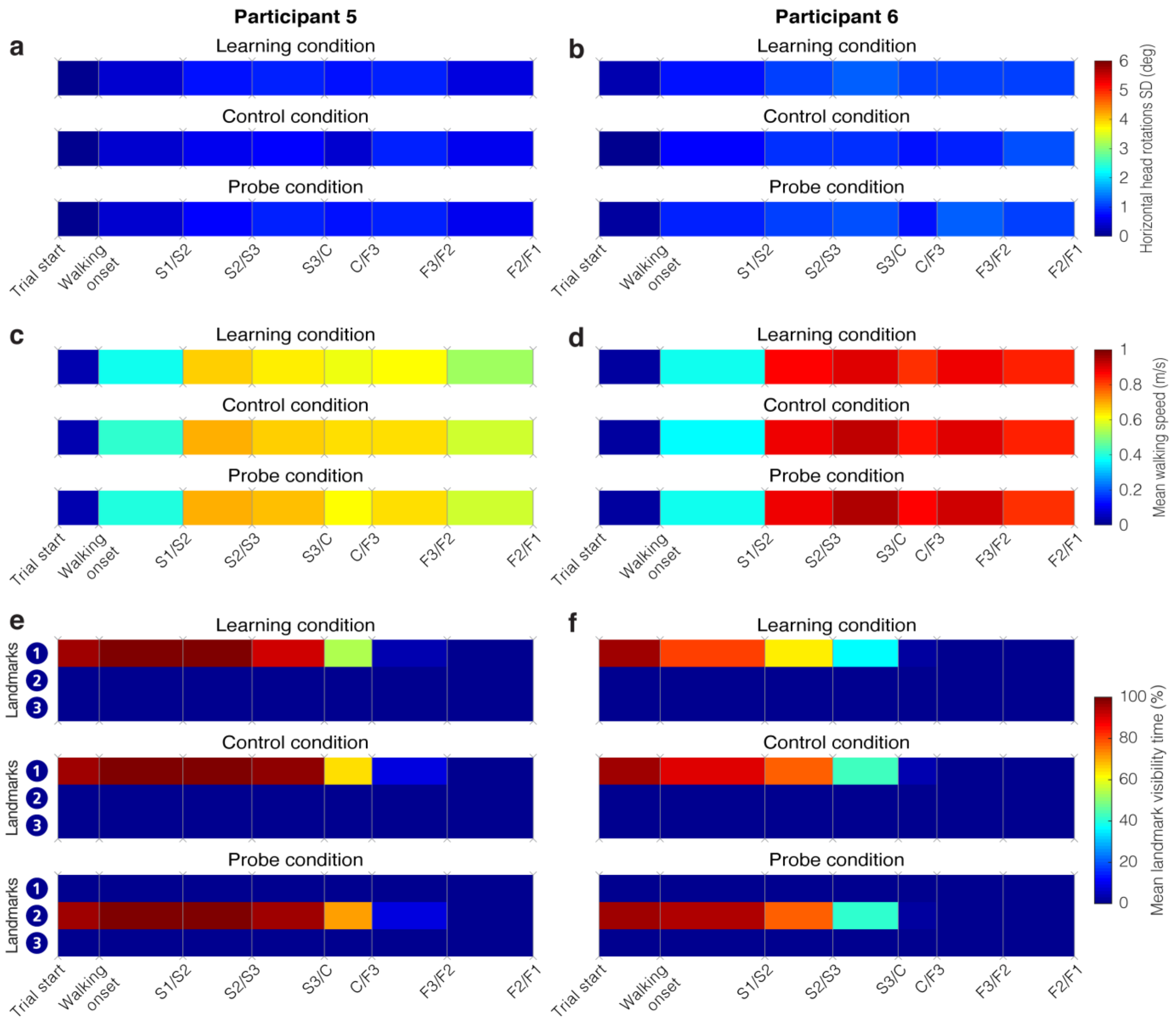

***Behavioral metrics – Walking speed, horizontal head rotations variability, and landmark visibility for the egocentric participants.*** **(a, c, e)** Participant 5. **(b, d, f)** Participant 6. For all plots, we divided each trial according to the same sequence of events: walking onset, followed by the first passage in the starting branch (S) then in the finish branch (F), being either the *goal* or the *error* branch. Events are horizontally spaced according to the median duration between each event. All plots represent data in the learning, the control and the probe conditions, averaged between separating events across all trials and blocks for each egocentric participant. **(a, b)** Average standard deviation of horizontal head rotations. **(c, d)** Average instantaneous walking speed. **(e, f)** Average landmark visibility. Color code corresponds to the percentage of time each landmark was visible at the screen.

#### **Supplementary Table 2.** ***3-way ANOVA on Landmark visibility***

| Factor | Degrees of freedom | F-statistic | p-value |
|---|---|---|---|
| **Main effects** | | | |
| **Condition** | (2;819) | 1.61 | 0.2 |
| **Zone** | (6;819) | 95.43 | < 0.00001 |
| **Landmark** | (2;819) | 263.50 | < 0.00001 |
| **2-way interaction effects** | | | |
| **Condition & Zone** | (12;819) | 0.13 | 1 |
| **Condition & Landmark** | (4;819) | 181.26 | < 0.00001 |
| **Zone & Landmark** | (12;819) | 23.54 | < 0.00001 |
| **3-way interaction effect** | | | |
| **Condition & Zone & Landmark** | (24;819) | 25.31 | < 0.00001 |

***Complete output of the 3-way ANOVA test on Landmark visibility.*** Effects and interactions for which the p-value was found below 0.01 were followed by post-hoc analyses involving pairwise t-tests between groups, corrected for multiple comparisons with Tukey's honest significant difference criterion method.

#### Supplementary Methods 1. *Motion Sickness questionnaire*

All participants answered a motion sickness questionnaire at the end of the experiment, adapted from Kennedy *et al.*, (1993). They had to rate the following symptoms as none, slight, moderate, or severe.

- General Discomfort
- Fatigue
- Eye Strain
- Headache
- Difficulty Focusing
- Salvation Increasing
- Sweating
- Nausea
- Difficulty concentrating
- Fullness of Head

#### Supplementary Methods 2. *BeMoBIL pipeline - Klug et al., (2018)*

##### *Bad channels detection*

EOG channels are excluded from the dataset at this step since they are likely to be considered as artifacts by the pipeline. Additionally, in preparation for the detection of bad channels, we removed portions of the continuous dataset that were not part of the actual trials to avoid taking them into account for the detection of abnormal channel behavior. To that purpose, 4 criteria are inspected:

- **Deviation criterion**. Find channels with extreme amplitudes. Extreme amplitudes are the sign of channels affected by large artifacts, suffering from poor contact with the scalp, etc...
- **Noisiness criterion**. Find channels with large high-frequency power. The signal of interest obeys a 1/f power function, therefore channels exhibiting abnormal power in the high-frequency band are likely to contain unusable signal.
- **Correlation criterion**. Find channels lacking correlation with any other channels. Because of scalp electrical conduction, channels should have a high level of correlation. When a channel has a signal very different from its neighbors, it is likely to be dysfunctional.
- **Predictability criterion**. Find channels lacking predictability by other channels. When group of channels are dysfunctional together, they might pass the previous criterion. A prediction drawn from other channels (not necessarily next to each other) should also respect a certain level of correlation with the original channel (again because of volume conduction).

We implemented this step with the *findNoisyChannels* function, taken from the PREP pipeline (Bigdely-Shamlo *et al.*, 2015). We set parameters numerical values according to default recommendations from Bigdely-Shamlo *et al.*, (2015).

##### *Noisy temporal segment detection*

The detection and removal of noisy temporal segments is particularly important for ICA decomposition as some periods affected by general artifacts may be interpreted as single ICs by the ICA algorithm (Delorme *et al.*, 2012; da Cruz *et al.*, 2018). Some portions of the continuous dataset were irrelevant to the scientific questions of this experiment (e.g.

disorientation phases). In our experiment, they are not necessarily noisier than other portions: for example, in disorientation phases, the participant walks but keeps his eyes closed and head relatively steady. This may be a particularly interesting situation to isolate artifacts generated by walking that will be similarly observed in the trials, in a messier situation. Hence, the possibility to use these portions was kept open (if they are not rejected by the noisiness detection), unlike for bad channels detection.

The BeMoBIL pipeline introduces an additional step before the actual detection of noisy temporal segments: artifacts are isolated and excluded from eye movements. The motivation for this lies in the fact that eye related artifacts yield large amplitude variations in the signal, hiding other artefacts to most metrics used for noisiness detection. Eye components are identified with ICA decomposition (AMICA, Palmer *et al.*, 2008) and automatic IC labelling (ICLabel, Pion-Tonachini *et al.*, 2019). To save computational time, we selected a smaller portion of the data to train the AMICA algorithm. This portion corresponded to the exploration phase plus the long baselines. We chose these phases because (1) they are equally defined for all participants; (2) their total duration (9 min) seemed suitable for training the ICA model in reasonable computational time; (3) they should provide examples of a variety of eye-related artifacts: blinks and slow eye movements when the participant finds himself immersed in the dark, large and fast eye movements, saccades to objects (paired with head movements) in the exploration phase. Eventually, we rely on the prediction given by the ICLabel algorithm to automatically identify eye components. Any IC for which the prediction exclusively exceeds the 'Eye' threshold is considered as an eye component. All eye components contribution to the channel-based dataset are removed with the *pop_subcomp* function from EEGLAB.

After this step, we band-pass filter the continuous stream between 1 and 40 Hz and then epoch the data into non-overlapping windows of 1 s. For each of these epochs, we compute 3 different quantities to evaluate their noisiness.

I. **Mean signal of the epoch** (averaged over channels and time). Large values point towards general impedance inflation or large artifacts affecting a large proportion of channels.
II. **Channel SD of epoch mean**. This is a simple measure of channel heterogeneity in the epoch. Large values indicate that some channels are affected by artifacts at an individual level in this period of time.

III. **Mahalanobian distance (MD) of epoch mean**. MD is a more robust estimation of channel heterogeneity than channel SD since it considers the variances and covariances between channels. Large values are also indicative of a noisy epoch.

Those quantities form a single score computed with a weighted sum giving more importance to the MD [w(I) = 1; w(II) = 1; w(III) = 2]. Then the epochs are sorted according to their score and the 15% highest scores are pinned for removal. Neighboring noisy epochs are merged to form blocks of rejection. Finally, each block is extended by 200 ms on both sides to account for artifact contamination of neighboring sections. We set parameters numerical values according to default recommendations from the authors of the pipeline.

#### Supplementary Figure 3. *Manual inspection of ICs labelling*

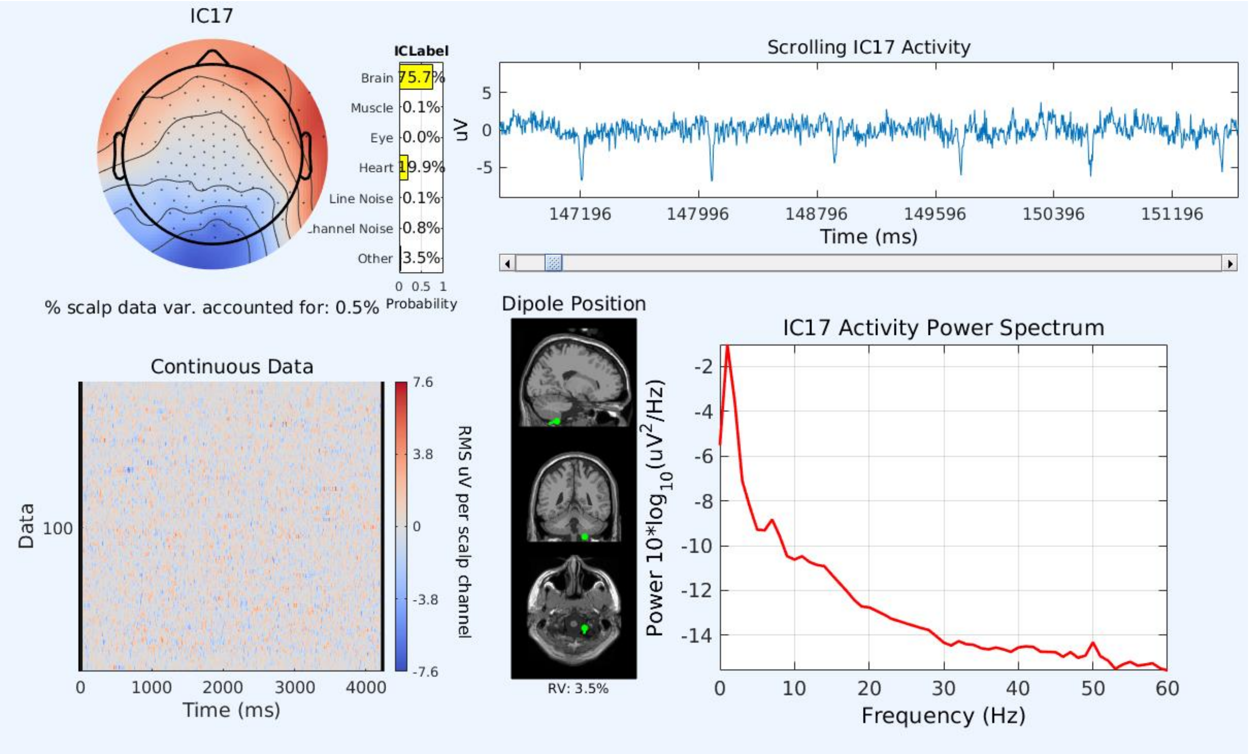

***Example of IC manual inspection during the assignment of IC labels.*** This IC from participant P09 would have been assigned to the Brain class without manual inspection. The IC was eventually labelled 'Heart' by the experimenter (heart beat clearly identifiable, gradient shape activation map and very deep ECD). This panel was extracted with the pop_prop_extended EEGLAB function.

#### Supplementary Methods 3. *Pipeline comparison*

##### *Pipeline evaluation*

To compare the performance of the BeMoBIL and the APP (da Cruz *et al.*, 2018) pipelines, we evaluated different metrics of the preprocessing process.

First, we inspected cleaning metrics: number of channels removed, percentage of data assigned to noisy temporal segments, percentage of brain labels among the retrieved ICs and the meaningfulness of these ICs quantified by the explained percentage of variance in the overall decomposition.

Second, as introduced by Delorme *et al.*, (2012), we chose 2 metrics to evaluate how well ICA achieved its independent decomposition objective.

- Mutual Information Reduction (MIR). It measures the difference between the mutual information in the original dataset (EEG channels) and the mutual information in the post-ICA dataset (ICs).
- Mean remaining Pairwise Mutual Information (PMI). The mutual information between a pair of ICs averaged over all pairs.

We employed non-parametric statistical tests to evaluate the pipelines against each other. When directly comparing the 2 main pipelines, we used the Wilcoxon signed rank test (WSRT) to assess the equality of medians (paired observations) and the Brown-Forsythe test (BFT) to assess the equality of spreads around median. We set the alpha level for significance at $p < 0.01$ for more conservative results.

##### *Comparison results*

We present the principal metrics for pipelines comparison on Supplementary Figure 3.

We first inspected the outcome of the cleaning steps where pipelines implemented different methods. There was no significant difference (WRST: $p = 0.12$) between the median number of channels removed by each pipeline (Supp. Fig. 3a), around 5 channels per subject. However, we found that the artefactual channel detection performed by the BeMoBIL pipeline was significantly more regular across subjects, with a distribution exhibiting a lower spread along the median than APP pipeline (BFT: $p = 0.005$). Moreover, the pipelines performed very different channel rejections: excluding the 3 subjects where the APP pipeline did not find any channels to reject, the median common percentage of rejection (expressed with respect to the pipeline rejecting less channels) is 33%. A detailed inspection of individual rejection criteria shows that the different implementation of similar criteria has a great impact: the deviation criterion as defined in the APP pipeline is significantly more sensitive than in the BeMoBIL pipeline (WRST: p = 2e-4, Supp. Fig. 3c) while we observe the opposite effect for the implemented correlation criterion (WRST: p = 3e-3, Supp. Fig. 3d).

On the contrary, we observed a significantly different cleaning behavior between pipelines at the bad temporal segment detection step (Supp. Fig. 3b). The APP pipeline rejected a median of about 5% of the total recording time per subject, against 17.5% for the BeMoBIL pipeline (WRST: p = 3e-5). The variability around this median is also significantly different between the pipelines (BFT: p = 2e-5) with a greater variability from APP than from BeMoBIL. A median of 80% of the time portions rejected by the APP pipeline were also rejected by the BeMoBIL pipeline.

Subsequently, we investigated the effect of each pipeline on the efficiency of ICA algorithm, measured by the reduction of mutual information. On a pairwise level, the mean remaining PMI (Supp. Fig. 3e) revealed a significantly greater performance of ICA after cleaning the data with the BeMoBIL pipeline than with the *APP* pipeline (WRST: p = 3e-5). We observe the same tendency at the scale of the global dataset (MIR, Supp. Fig. 3f) but with no significant difference (WRST: p = 0.14).

In conclusion, the BeMoBIL pipeline demonstrated more robustness and conservativeness than the APP pipeline in the artefact detection steps. The underlying adaptability proposed by APP pipeline is not advisable as its performance proved to be very inconsistent across participants. When released, this pipeline had not been tested on mobile data (da Cruz *et al.*, 2018) and the particularities of such recordings, prone to withhold a large spectrum of unusual artifacts (related to gait, large head movements, cable pulling, …) had not been considered. More importantly, according to the mutual information reduction metrics, the BeMoBIL pipeline provided a better preparation for the ICA decomposition than APP pipeline, enabling a greater independence between the resulting ICs.

#### **Supplementary Figure 4.** ***Pipeline comparison results***

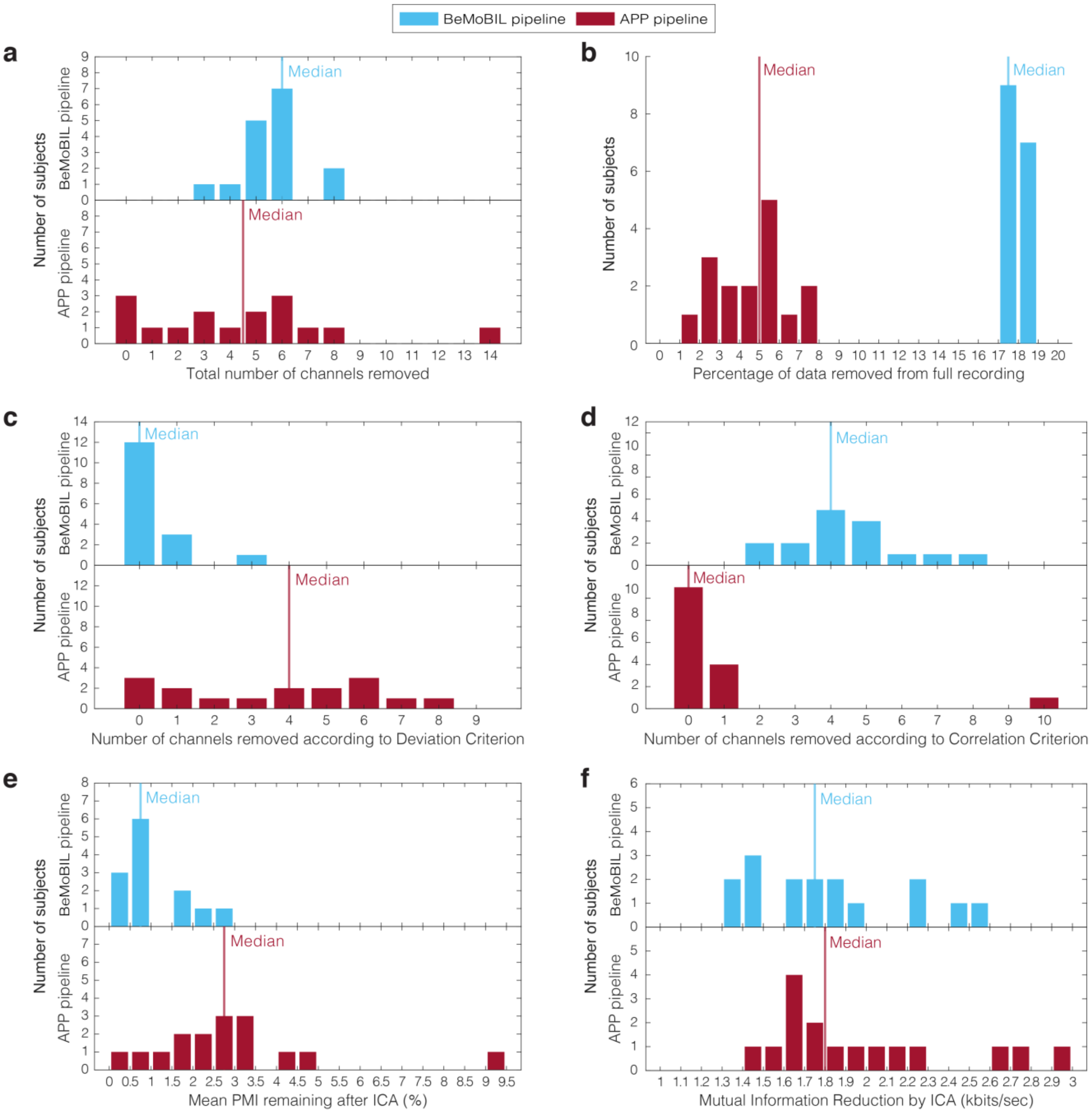

***Pipeline comparison metrics.*** **(a)** ***Artefactual channels outcome.*** Histogram plots showing the distribution of number of artefactual channels identified by each pipeline for each subject (N=16). WSR test (BeMoBIL-APP): U = 77.5; p = 0.12. BF test: F(1;30) = 9.10; p = 0.005. subfigures **(c)** and **(d)** show the detail of this identification depending on the criteria used: the *deviation criterion* **(c)** and the *correlation criterion* **(d)** are implemented differently in the pipelines. **(b)** ***Temporal artefacts detection outcome.*** Histogram plots showing the distribution of the percentage (with respect to total recording time) of time segments detected as artefactual by each pipeline for each subject (N=16). WSR test (BeMoBIL-APP): U = 136; p = 3e-5. BF test: F(1;30) = 24.28; p = 2e-5. **(e) *Remaining Pairwise Mutual Information after ICA decomposition.*** Histogram plots showing the distribution of the mean remaining PMI for the ICA decomposition after each pipeline. For each subject, we first computed the PMI of all pairs (separately in the channel and the component spaces) and then averaged over all pairs in each space. Remaining PMI is the ratio of ICs mean over channels mean, presented as a percentage. WSR test (BeMoBIL-APP): U = 0; p = 3e-5. BF test: F(1;30) = 2.37; p = 0.13. **(f) *Mutual Information Reduction achieved by ICA decomposition.*** Histogram plots showing the distribution of the MIR for the ICA decomposition after each pipeline. MIR is the difference of global mutual information contained in the dataset between the IC representation and the channel representation. WSR test (BeMoBIL-APP): U = 39; p = 0;14. BF test: F(1;30) = 0.19; p = 0.67.

### Supplementary Methods 4. *Choice of the clustering design parameters*

#### *Designs definition - Parameters inspected*

To inspect the influence of k-means clustering algorithm parameters, we compared four sets of parameters: number of formed clusters could alternatively be 50 or 60 and threshold for outliers was either 3 or 4 SD. Setting the number of clusters below the number of ICs per participant is common practice (Luu *et al.,* 2017b; Gramann *et al.,* 2018; Nordin *et al.,* 2019) as there is no guarantee for the activity associated with a cortical region to be represented by a unique IC. Additionally, we evaluated the best clustering solution with respect to four different possible RSC coordinates as ROI. We took the first location (RSC1, [0,-45,10]) from Gramann *et al.* (2018) , the second one (RSC2, [0,-56,9]) from Lin *et al.* (2015), the third one (RSC3, [0,-47,7]) from Shine *et al.* (2016) and we chose the last one close to the anatomical region BA30 (RSC4, [0,-55,15]). We set the first coordinate (x) to 0 because we did not have any expectation for lateralization. Coordinates are expressed in MNI format.

#### *Metrics for ranking solutions within design*

For each design, we scored the clustering solutions following the procedure described in Gramann *et al.* (2018). For each of the 10000 clustering solutions, we first identified the cluster whose centroid was closest to the target ROI. Then, we inspected it using 6 metrics: (1) number of participants represented in the cluster, (2) ratio of ICs per participant, (3) cluster spread (normalized to the number of ICs in the cluster), (4) mean RV, (5) distance between cluster centroid and ROI coordinates and (6) Mahalanobis distance to the median of the solutions. We combined these metrics (after normalization) in a single score using a weighted sum [w1=2, w2=-3, w3=-1, w4=-1, w5=-3, w6=-1] and eventually clustering solutions were ranked according to their score.

#### *Metrics for design comparison*

We compared designs (*i.e.* set of parameters) based on the evaluating metrics for their highest rank solution and the stability of these metrics across the 11 best ranking solutions. We computed a *design score* out of the evaluation metrics for the highest rank solution, normalized across the 16 designs, with the same weights (supplementary Equation 1). For each metric, we also assessed how stable the value of the highest rank solution was among the 10 following solutions found with the same design parameters, using the variability index in supplementary

Equation 2. We summarized the overall variability with a single score computed from the weighted average of the variability for each measure (supplementary Equation 3). Finally, we selected the design with 50 clusters, 3 SD as threshold for outliers and RSC4 set of coordinates for target ROI.

***Supplementary Equation 1:***

Summary solution score.

$$\boldsymbol{SCORE(D_i) = \sum_{j=1}^{6} w_j * M_j^{normalized}}$$

$$\boldsymbol{with\ D_i = design\ i, M_j^{normalized} = measure\ j\ normalized\ across\ designs\ and}$$
$$\boldsymbol{w_j = weigth\ for\ measure\ j}$$

***Supplementary Equation 2:***

Variability index assessing the stability of a best rank solution measure across the following best ranked solutions.

$$\boldsymbol{VAR_{11}(M_i) = 100 * \frac{mean_{j=2\rightarrow 11}(|M_i(Sol_j) - M_i(Sol_1)|)}{M_i(Sol_1)}}$$

$$\boldsymbol{with\ M_i = measure\ i\ and\ Sol_j = solution\ of\ rank\ j}$$

***Supplementary Equation 3:***

Summary variability index.

$$\boldsymbol{VAR(D_i) = \sum_{j=1}^{6}|w_j| * VAR_{11}(M_j) \Big/ \sum_{j=1}^{6}|w_j|}$$

$$\boldsymbol{with\ D_i = design\ i, M_j = measure\ j\ and\ w_j = weigth\ for\ measure\ j}$$

##### *Choice of clustering parameters*

We present the scores comparing the different clustering parameters in supplementary Table 3. Increasing the SD threshold (σ in the table) for outliers unequivocally yield worse solutions for this dataset, mainly due to the fact that the 1:1 ratio between number of ICs and number of participants is lost. Within designs with 3 SD threshold, RSC1 set of coordinates outputted very singular solutions, with fewer participants than other ROIs and associated with a high variability score indicating that those solutions were not representative of the pool of best ranks solutions for these designs. The solutions coming from designs with other ROIs were generally more stable. RSC2, RSC3 and RSC4 solutions retrieved almost identical clusters but the set of coordinates consistently closest to the centroid of this cluster was RSC4. We therefore opted for a design with this parameter. The remaining 2 designs (50 or 60 clusters with 3SD and RSC4) were associated to similar scores (highest ones amongst the 16 designs) and variability scores (low variability in each case). We eventually chose the 50 clusters design to favor the analysis of bigger clusters, potentially regrouping ICs from a larger share of participants and therefore more representative of our population. Choosing the RSC coordinates without any reference, has to be put in perspective with the high variability across literature of RSC functional location (Epstein, 2008) and the poorer spatial resolution of source localization with respect to fMRI scans.

**Supplementary Table 3.** ***Results of the cluster design comparison***

| Design | | | Measures | | | | | | | | | | | | | | | |
|---|---|---|---|---|---|---|---|---|---|---|---|---|---|---|---|---|---|---|
| Nb. of clusters | σ | ROI | Nb of Participants | | Mean IC/Part. | | Cluster spread | | Mean RV (%) | | Centroid-ROI distance | | Mahalanobis distance | | SCORE | | VAR | |
| | | | BEST | VAR11 | BEST | VAR11 | BEST | VAR11 | BEST | VAR11 | BEST | VAR11 | BEST | VAR11 | value | rank | value | rank |
| 60 | 3 | RSC1 | 5 | 66,0 | 1 | 0,0 | 171,6 | 20,4 | 3,29 | 10,6 | 8,05 | 48,9 | 29,63 | 43,8 | -3,681 | 3 | 32,145 | 16 |
| 60 | 3 | RSC2 | 12 | 2,5 | 1 | 1,5 | 215,0 | 1,7 | 4,36 | 1,3 | 18,11 | 1,6 | 9,75 | 5,9 | -3,744 | 4 | 2,117 | 1 |
| 60 | 3 | RSC3 | 12 | 18,3 | 1 | 0,8 | 215,0 | 6,6 | 4,36 | 7,8 | 18,86 | 13,8 | 9,16 | 92,0 | -3,844 | 7 | 16,990 | 13 |
| 60 | 3 | RSC4 | 12 | 2,5 | 1 | 1,5 | 215,5 | 1,7 | 4,11 | 6,7 | 13,23 | 1,9 | 11,66 | 15,5 | -2,980 | 1 | 3,575 | 2 |
| 50 | 3 | RSC1 | 9 | 27,8 | 1 | 3,3 | 213,3 | 2,2 | 3,58 | 22,8 | 12,86 | 23,7 | 25,22 | 40,9 | -3,798 | 6 | 18,404 | 14 |
| 50 | 3 | RSC2 | 12 | 0,8 | 1 | 7,4 | 215,0 | 2,2 | 4,36 | 2,2 | 18,11 | 2,4 | 10,46 | 10,7 | -3,768 | 5 | 4,207 | 5 |
| 50 | 3 | RSC3 | 12 | 2,5 | 1 | 5,0 | 215,0 | 1,5 | 4,36 | 2,5 | 18,86 | 3,3 | 16,83 | 18,2 | -4,103 | 8 | 4,745 | 7 |
| 50 | 3 | RSC4 | 12 | 0,8 | 1 | 7,4 | 215,0 | 2,2 | 4,36 | 2,2 | 13,10 | 1,3 | 10,84 | 10,1 | -2,985 | 2 | 3,855 | 4 |
| 60 | 4 | RSC1 | 12 | 5,8 | 1,083 | 2,2 | 246,5 | 7,4 | 3,87 | 13,5 | 14,19 | 11,1 | 11,76 | 9,8 | -4,828 | 10 | 7,480 | 10 |
| 60 | 4 | RSC2 | 13 | 7,7 | 1,077 | 0,6 | 224,8 | 2,4 | 4,52 | 1,2 | 17,98 | 1,7 | 21,47 | 21,0 | -5,512 | 13 | 4,255 | 6 |
| 60 | 4 | RSC3 | 12 | 4,2 | 1,083 | 2,1 | 246,5 | 8,0 | 3,87 | 14,9 | 16,84 | 7,4 | 11,98 | 7,6 | -5,258 | 12 | 6,109 | 8 |
| 60 | 4 | RSC4 | 13 | 7,7 | 1,077 | 0,6 | 224,8 | 2,4 | 4,52 | 1,2 | 12,98 | 0,0 | 19,47 | 21,2 | -4,649 | 9 | 3,811 | 3 |
| 50 | 4 | RSC1 | 11 | 11,8 | 1,091 | 5,0 | 238,9 | 17,1 | 3,75 | 16,4 | 13,63 | 15,5 | 18,42 | 36,5 | -5,240 | 11 | 14,108 | 12 |
| 50 | 4 | RSC2 | 13 | 13,1 | 1,154 | 4,2 | 234,0 | 8,7 | 4,49 | 4,6 | 18,46 | 7,6 | 24,33 | 16,3 | -7,214 | 16 | 8,282 | 11 |
| 50 | 4 | RSC3 | 8 | 40,0 | 1,125 | 4,8 | 255,0 | 20,6 | 4,18 | 12,3 | 11,02 | 44,4 | 16,58 | 41,8 | -6,130 | 15 | 27,483 | 15 |
| 50 | 4 | RSC4 | 13 | 9,2 | 1,154 | 6,4 | 234,0 | 11,4 | 4,49 | 4,0 | 13,19 | 2,7 | 12,40 | 13,3 | -5,974 | 14 | 6,761 | 9 |

***Measures for the best solution outputted by each clustering design.*** The first 3 columns introduce the clustering design parameters (see "Designs definition - Parameters inspected" section), namely the target number of clusters, the outliers' threshold and the ROI coordinates. The middle 6 columns show the evaluation of each design along a single metric, as presented in the "Metrics for ranking solutions within design". *BEST* sub-column corresponds to the metric value associated to the best of the 10000 solutions for the given design parameters. *VAR11* sub-column corresponds to a variability index assessing the stability of a best rank solution measure across the following 10 best ranked solutions (see Supplementary Equation 2). The last 2 columns present summary scores aggregating the weighted contribution of all metrics to the ranking of design parameters. *SCORE* column corresponds to the summary solution score (see Supplementary Equation 1) and *VAR* column corresponds to the summary variability index (see Supplementary Equation 3). The *rank* sub-column evaluates the ordering of clustering parameters according to the given summary column (ranked from 1 to 16 with 1 associated to the best performance).

#### Supplementary Table 4. *Brain cluster selection*

| Clust. ID | Nb. allo. part. | Nb allo. ICs | Mean Position | | | Mean dist. to centroid (mm) | Mean RV | STD RV | Talairach Client: Closest Gray Matter region | | | | | | Kept for later analysis |
|---|---|---|---|---|---|---|---|---|---|---|---|---|---|---|---|
| | | | x | y | z | | | | Level 1 | Level 2 | Level 3 | Level 4 | Level 5 | Range (mm) | |
| 1 | 10 (12) | 10 (12) | 7,67 | -46,84 | 24,47 | 13,6 | 4,4% | 2,4% | Right Cerebrum | Limbic Lobe | Posterior Cingulate | Gray Matter | Brodmann area 23 | 2 | YES |
| 2 | 11 (12) | 21 (22) | 15,19 | -81,61 | 35,00 | 9,8 | 6,4% | 2,4% | Right Cerebrum | Occipital Lobe | Cuneus | Gray Matter | Brodmann area 19 | 0 | YES |
| 3 | 9 (11) | 13 (15) | 38,96 | -50,54 | 32,91 | 10,9 | 7,3% | 4,1% | Right Cerebrum | Parietal Lobe | Supramarginal Gyrus | Gray Matter | Brodmann area 40 | 4 | YES |
| 4 | 11 (12) | 14 (15) | -2,38 | 9,63 | 21,89 | 15,6 | 3,5% | 3,1% | Left Cerebrum | Limbic Lobe | Anterior Cingulate | Gray Matter | Brodmann area 33 | 2 | YES |
| 5 | 11 (13) | 15 (17) | 33,26 | -9,74 | 52,04 | 14,6 | 7,1% | 5,1% | Right Cerebrum | Frontal Lobe | Precentral Gyrus | Gray Matter | Brodmann area 6 | 1 | YES |
| 6 | 10 (11) | 12 (13) | -37,34 | -27,53 | 48,87 | 11,7 | 6,5% | 4,4% | Left Cerebrum | Parietal Lobe | Postcentral Gyrus | Gray Matter | Brodmann area 3 | 0 | YES |
| 7 | 8 (9) | 10 (11) | -10,50 | -55,98 | 39,15 | 10,7 | 5,4% | 2,1% | Left Cerebrum | Parietal Lobe | Precuneus | Gray Matter | Brodmann area 7 | 2 | NO |
| 8 | 8 (8) | 8 (8) | 26,04 | 29,69 | 26,20 | 14,3 | 5,2% | 2,0% | Right Cerebrum | Frontal Lobe | Middle Frontal Gyrus | Gray Matter | Brodmann area 9 | 4 | NO |
| 9 | 8 (8) | 11 (11) | 1,33 | -30,51 | 60,23 | 11,5 | 7,1% | 5,1% | Right Cerebrum | Frontal Lobe | Paracentral Lobule | Gray Matter | Brodmann area 6 | 3 | NO |
| 10 | 8 (10) | 11 (13) | 4,28 | -17,39 | 0,35 | 15,4 | 5,2% | 4,2% | Right Cerebrum | Sub-lobar | Thalamus | Gray Matter | * | 0 | NO |
| 11 | 7 (8) | 7 (8) | -30,74 | -58,97 | 23,64 | 13,1 | 6,9% | 3,0% | Left Cerebrum | Temporal Lobe | Middle Temporal Gyrus | Gray Matter | Brodmann area 39 | 3 | NO |
| 12 | 5 (7) | 5 (7) | 55,87 | -20,08 | -27,43 | 14,4 | 9,6% | 1,9% | Right Cerebrum | Temporal Lobe | Fusiform Gyrus | Gray Matter | Brodmann area 20 | 1 | NO |

***Selection among the 12 Brain clusters.*** For the rest of the analysis, we chose to keep only the clusters containing ICs from at least 9 out of the 14 allocentric participants (~65%). This table presents all 12 brain clusters retrieved from the clustering procedure described above. In the order of the columns from left to right:

- Cluster ID (clusters 1 to 6 are presented in Fig. 7 & Fig. 8 in the main document),
- Number of allocentric participants (*resp.*, in parenthesis, total number of participants) presenting at least one IC in the cluster,
- Number of ICs accumulated by allocentric (*resp.*, in parenthesis, all) participants in the cluster,
- Mean position of the cluster centroid in TAL coordinates,
- Mean distance of the cluster ICs to the centroid (mm),

- Mean residual variance (%),
- Standard deviation to the residual variance (%),
- Closest gray matter region as located by the Talairach Client (Lancaster *et al.*, 2000),
- Decision to keep the clusters for the rest of the analysis. Cluster 1 to 6 were kept.
